# A multi-omics approach to visualize early neuronal differentiation in 4D

**DOI:** 10.1101/2022.02.01.478732

**Authors:** Athina Samara, Mari Spildrejorde, Ankush Sharma, Martin Falck, Magnus Leithaug, Stefania Modafferi, Pål Marius Bjørnstad, Ganesh Acharya, Kristina Gervin, Robert Lyle, Ragnhild Eskeland

**Affiliations:** Division of Clinical Paediatrics, Department of Women’s and Children’s Health, Karolinska Institutet, Sweden; Astrid Lindgren Children′s Hospital Karolinska University Hospital, Stockholm,Sweden; PharmaTox Strategic Research Initiative, Faculty of Mathematics and Natural Sciences, University of Oslo, Norway; Department of Medical Genetics, Oslo University Hospital and University of Oslo, Norway; Institute of Clinical Medicine, Faculty of Medicine, University of Oslo, Oslo, Norway; Department of Informatics, University of Oslo, Norway; Institute of Basic Medical Sciences, University of Oslo, Oslo, Norway; Department of Biosciences, University of Oslo, Norway; Division of Obstetrics and Gynecology, Department of Clinical Science, Intervention and Technology (CLINTEC), Karolinska Institutet, Alfred Nobels Allé 8, SE-14152, Stockholm, Sweden; Center for Fetal Medicine, Karolinska University Hospital Huddinge, SE-14186 Stockholm, Sweden; Pharmacoepidemiology and Drug Safety Research Group, Department of Pharmacy, School of Pharmacy, University of Oslo, Norway; Division of Clinical Neuroscience, Department of Research and Innovation, Oslo University Hospital, Oslo, Norway; Centre for Fertility and Health, Norwegian Institute of Public Health, Oslo, Norway

**Keywords:** Single-cell RNA-seq, scATAC-seq, human embryonic stem cells, neuronal differentiation, DNA methylation, telencephalic signatures

## Abstract

Neuronal differentiation of pluripotent stem cells is an established method to study physiology, disease and medication safety. However, the sequence of events in human neuronal differentiation and the ability of *in vitro* models to recapitulate early brain development are poorly understood. We developed a protocol optimized for the study of early human brain development and neuropharmacological applications. We comprehensively characterized gene expression and epigenetic profiles at four timepoints, as the cells differentiate from embryonic stem cells towards a heterogenous population of progenitors, immature and mature neurons bearing telencephalic signatures. A multi-omics roadmap of neuronal differentiation, combined with searchable interactive gene analysis tools, allows for extensive exploration of early neuronal development and the effect of medications.

**Graphical Abstract:** 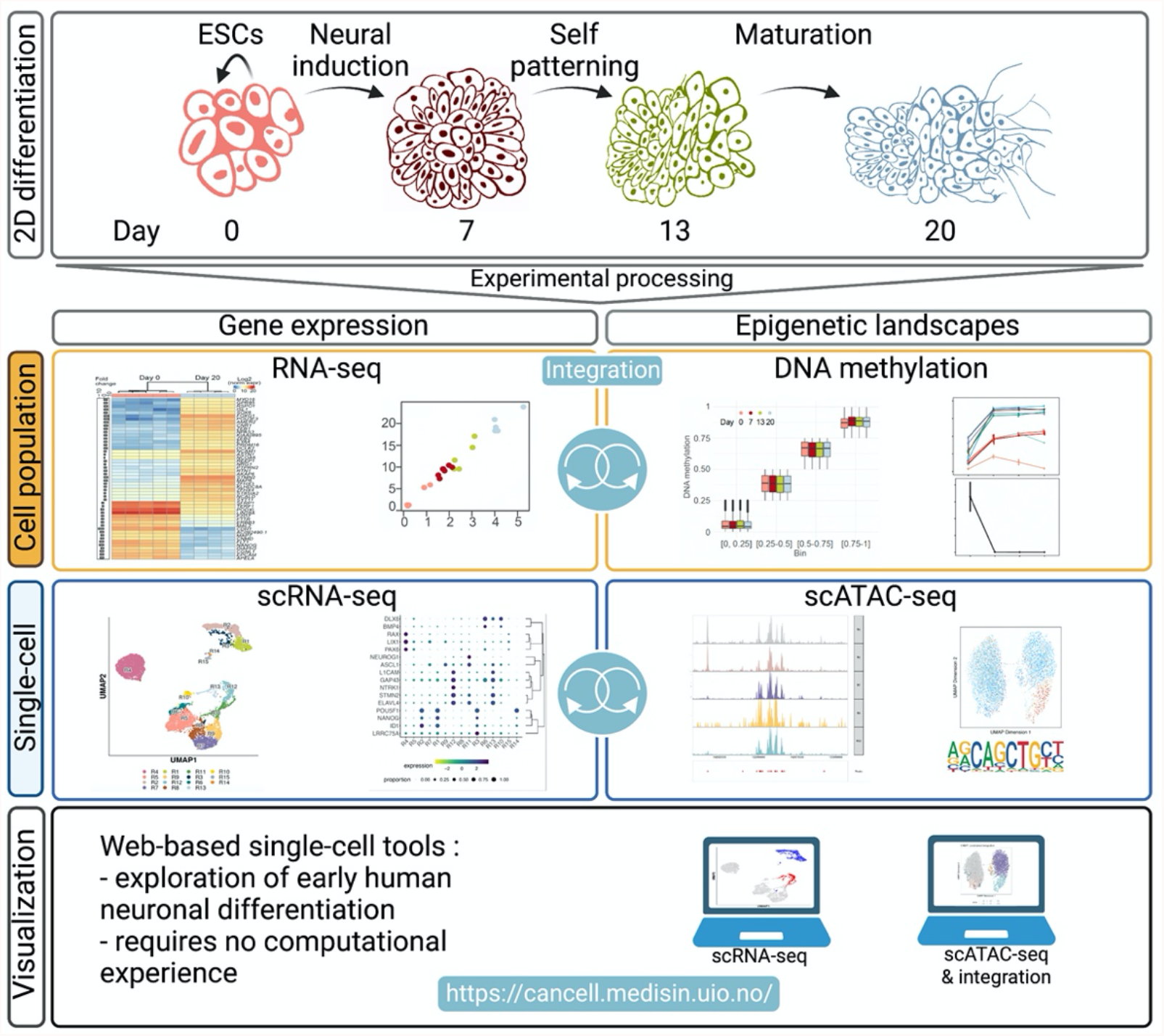

**Highlights:**

- Multi-omics charting a new neuronal differentiation protocol for human ES cells
- Single-cell analyses reveals marker genes during neuronal differentiation
- Identified transcriptional waves similar to early human brain development
- Searchable tools to visualize single-cell gene expression and chromatin state

**In Brief:** We have developed a novel protocol for human embryonic stem cells to study neural induction and early neuronal differentiation. Multi-omics analyses uncovered cell populations, genes and transcriptional waves defining cell fate commitment. We comprehensively describe epigenetic landscapes and gene expression and provide searchable analysis tools for exploration of the data.

## Introduction

Neuronal differentiation of pluripotent stem cells (PSCs) is an established method used to study early development, physiology, disease and neurotoxicity (Riemens et al., 2018). However, there is a need for robust protocols that systemically characterize cells at intermediate differentiation timepoints. These types of *in vitro* studies should offer the cell-type resolution necessary to characterize developmental trajectories. Improving the understanding of a protocol’s ability to recapitulate early brain development will aid future studies and increase applicability.

The role of epigenetic regulation on the establishment and maintenance of cellular identity during early neuronal differentiation processes is not well understood (Sun et al., 2021; Yao et al., 2016). Therefore, in-depth analyses describing epigenetic landscapes and the complex interplay with gene expression are required. Moreover, mapping derivative cells and their development- or region-specific transcriptional and epigenetic landscapes is fundamental for investigating disease mechanisms and for therapeutic interventions.

In this study, we used a multi-omics approach to construct a molecular timeline of early human neuronal differentiation. We used a novel 2D neuronal differentiation protocol using dual SMAD/WNT signalling inhibitors LDN193189, SB431542 and XAV939 (LSX) for neural induction of human embryonic stem cells (hESCs) (Cakir et al., 2019; Chavali et al., 2020; Major et al., 2016; Ohashi et al., 2018; Tchieu et al., 2017). The neuronal progenitors were allowed to self-pattern and mature towards a heterogenous population of immature and mature neurons bearing telencephalic signatures. We performed RNA-seq, global DNA methylation, single-cell RNA-seq and ATAC-seq data integration across timepoints (4D analysis), to correlate the expression of transcription factors with time- and population-specific chromatin states in hESCs, and during differentiation. This integration of comprehensive multi-omics data enabled the characterization of both the transcriptional and epigenetic landscapes in this model of early fate commitment. We provide access to single-cell data in user-friendly, interactive web applications that enable visualization of gene cluster regulation during the neuronal differentiation protocol.

## Results

### Initial validation of the neuronal differentiation protocol

The coating conditions and cell numbers were optimized to permit high cell contact, proliferation, and viability. Thus, based on confluency, morphology and viability, we analyzed the hESCs (Day 0) and derivative cell populations at three timepoints. We defined the end of the neural induction phase (Stage I) at Day 7, the end of the self-patterning phase (Stage II) at Day 13 and at the end of the maturation phase (Stage III) at Day 20 (**Fig. 1A**). For the neural induction of undifferentiated HS360 hESCs (Main et al., 2020; Ström et al., 2010), we used LDN193189, SB431542 and XAV939 (LSX). This LSX cocktail antagonizes the BMP, TGFβ and WNT signalling pathways to drive cells to anterior neuroectoderm (Cakir et al., 2019; Major et al., 2016; Ohashi et al., 2018; Tchieu et al., 2017). By the end of Stage I, neural induction morphogenetic events shape cells into thickened neural rosettes, whereas at Stage II, cells self-pattern before the Stage III FGF2/EGF maturation phase (**Fig. 1B**). In the absence of inhibitors at the self-patterning stage II, the cells retain their anterior forebrain identity and proceed to maturation, as shown by the ddPCR results (**Fig. 1C**).

**Figure 1.**
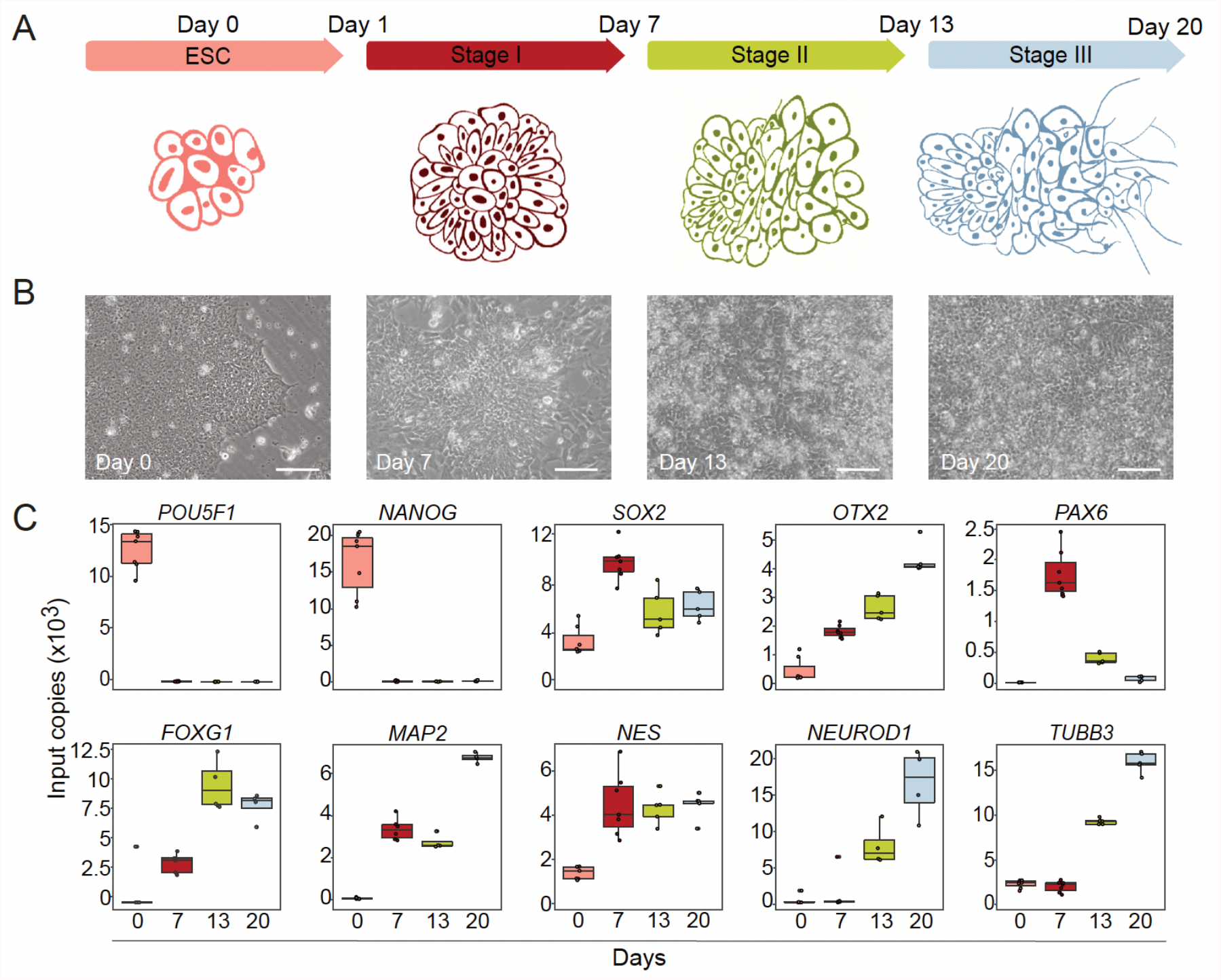
2D protocol with neural induction followed by self-patterning and maturation. A) Schematic illustration of the 20 Day timeline of the neuronal differentiation protocol from hESCs. B) Representative 20x brightfield phase contrast images of hESCs at Days 0, 7, 13 and 20 (scale bar 100 μm). C) ddPCR results from 4-6 replicates of mRNA expression of selected marker genes from Days 0, 7, 13 and 20.

The expression of the pluripotency markers *POU5F1* and *NANOG* decreased significantly after neural induction (p < 0.00001). Expression of the early neural markers *SOX2* and *NES* increased and stabilized at Day 7, whereas *PAX6* expression peaked at Day 7 before decreasing significantly at Days 13 and 20 (p < 0.0001). The expression of the transcription factor (TF) *OTX2*, which regulates neurogenesis and antagonizes ground state pluripotency, the late onset pan-neuronal marker *TUBB3*, and also *MAP2 and FOXG1* increased as cells differentiated. Immunofluorescence imaging showed protein expression and localization of OCT4, OTX2, SOX2, PAX6, NES and TUBB3 (**Fig. S1**).

### Identification of heterogenous populations of progenitors, mature and immature neurons with telencephalic signatures

To characterize the gene expression signatures, composition, differentiation pathway trajectories and the maturation level of the cell types derived, we performed single-cell RNA-seq (scRNA-seq) analyses at Days 0, 7, 13 and 20 (**Figs. 2, S2, S3** and **Table S1**). The scRNA-seq data can be visualized in the open access webtool “hESC Neuronal Differentiation scRNA-seq” (**hESCNeuroDiffscRNA**) where expression of genes can be explored per cell, cluster and timepoint (**Star methods**). A total of 9,337 cells were projected in UMAPs, 1,900 Day 0 cells, 2,368 Day 7cells, 2,045 Day 13 cells and 3,024 Day 20 cells). (**Fig. 2A** and **hESCNeuroDiffscRNA**, cell information tab, orig.ident).

**Figure 2.**
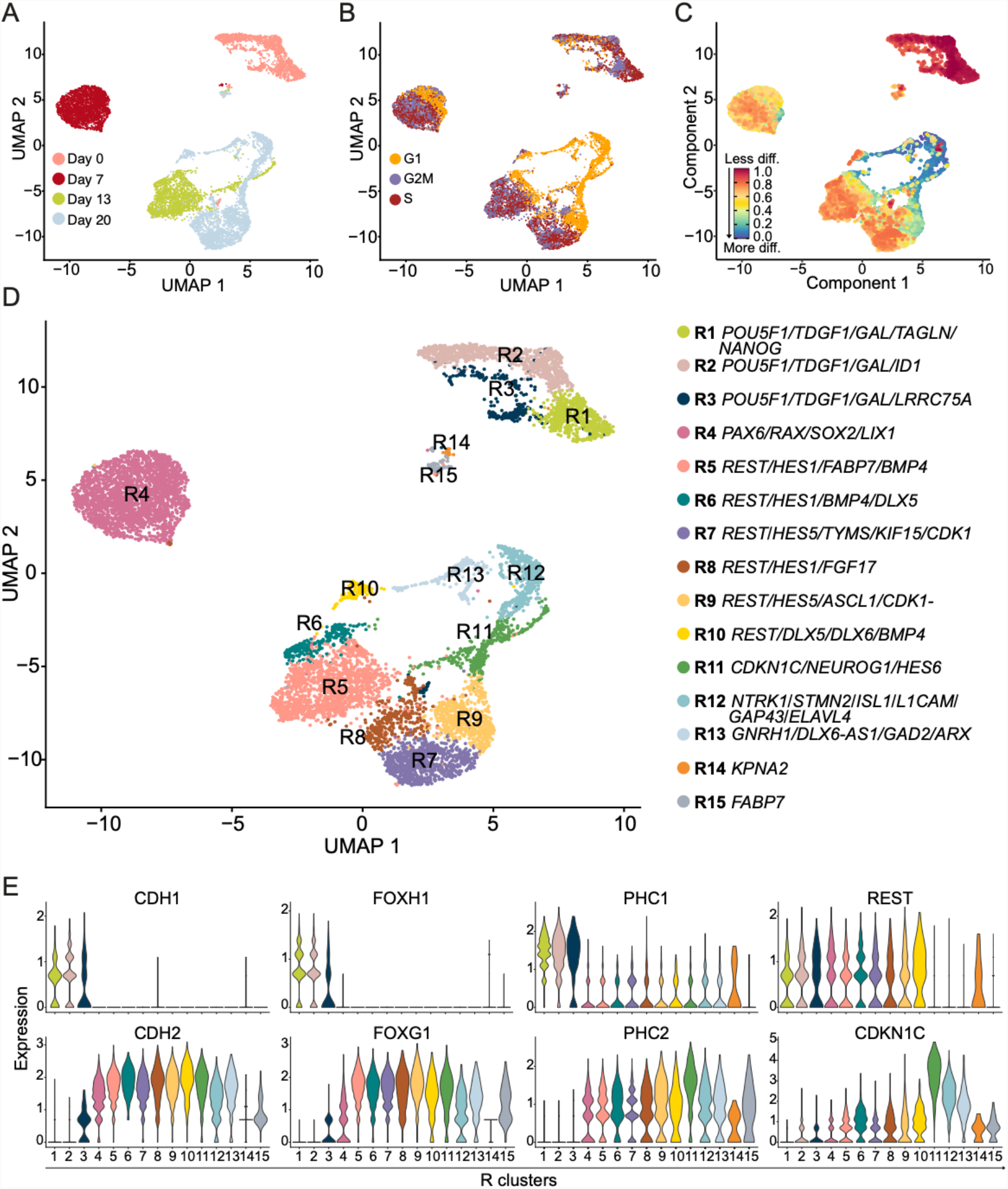
Identification of cell populations during neuronal differentiation of hESCs. A) UMAP representing single-cell RNA-seq clusters per time-point. B) UMAP of cell cycle analysis showing all cells analyzed and coloured by assigned cell cycle phase. C) The inferred neuronal differentiation trajectory using CytoTRACE, where less differentiated cells are shown in red and more differentiated cells are shown in blue. D) The UMAP representing cell clusters R1 to R15 with corresponding gene annotations mapped at resolution 0.55. The clusters are indicated by different colours and gene annotations per cluster are given after each colour corresponding bullet below the UMAP. E) Violin plots representing gene expression levels and distribution in clusters R1 to R15 for selected genes.

### Inferring quantitative analysis of cell cycle phase

A hallmark of neuronal development involves major alterations in G1- and S-phase duration. G1-phase lengthening is associated with the transition to a more differentiated cell type, while S-phase duration is linked to progenitor cell expansion (Arai et al., 2011). The cell cycle-specific gene trajectories showed a transition from 15.4 % to 54.1% cells in G1 phase from Day 0 to 20 (**Fig. 2B, S2D, Table S2** and **hESCNeuroDiffscRNA**). This is consistent with previous studies showing that the maintenance of pluripotency, proliferation and differentiation of rapidly proliferating PSCs, neural stem cells and progenitor cells are regulated by the cell cycle (Becker et al., 2006; Boward et al., 2016; Liu et al., 2019; Soufi and Dalton, 2016). The cell cycle regulator *CDK1* was expressed in 60% of the cells at Day 0, 45% at Day 7, 53% at Day 13, and reduced to 33% at Day 20 (**Fig. S2E**, % from the **hESCNeuroDiffscRNA**). CytoTRACE results confirmed that differentiation is consistent with the cell cycle phase inferred trajectory. As indicated by the higher CytoTRACE scores, cell potency gradually decreased from Day 0 to 20 (**Fig. 2C**), confirming the cell cycle phase prediction.

### Development and differentiation markers used for cluster resolution and annotation

The four timepoints were resolved into 15 clusters (R1-R15, **Fig. 2D**). Corresponding cell numbers per cluster and cells per timepoint per cluster are shown (**Table S3)**. The top ten most highly expressed genes for each cluster are plotted in a heatmap (**Fig. 3**), including many developmentally regulated TFs. Among these genes, *POU5F1, TDGF1, GAL, LRRC75A, RAX, LIX1, TYMS, HES1, HES5, HES6, FGF17, DLX5, DLX6, GAP43, STMN2* and *GNRH1* were used for R1-R13 cluster annotation. For R14, consisting of 27 Day 0 cells, we used *KPNA2*, a gene associated with the localization of OCT4 (Li et al., 2008). For R15, a pool of 90 cells from Days 7, 13 and 20, we used *FABP7*, which is expressed in NSCs during development (Kurtz et al., 1994).

**Figure 3.**
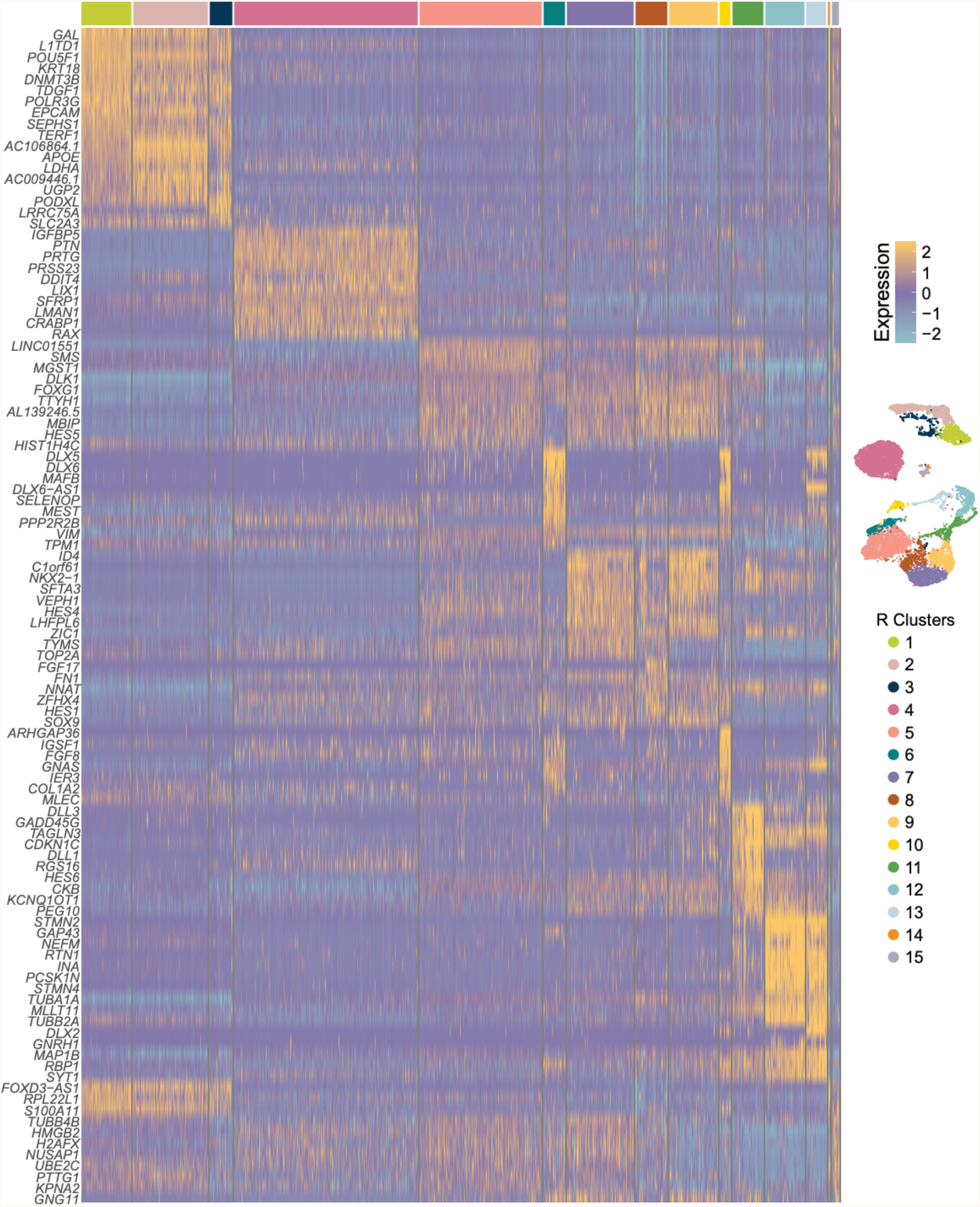
Cluster analysis of differential gene expression. Heatmap of the top ten most highly expressed genes in all clusters. Rows represents single genes and column represents single cells. Cell clusters are ordered sequentially and coloured according to the UMAP annotation shown to the right (yellow represents high expression and blue low expression).

### Characterizing the unsynchronized hESC population

We identified three distinct Day 0 clusters (R1-3) where all cells expressed *POU5F1*, verifying their pluripotency. TFs essential in establishing and maintaining pluripotency (i.e., *GAL, TDGF1, ID1, FOXH1* and *SOX2)* were highly expressed in clusters R1-3 (observe in **hESCNeuroDiffscRNA**). R3 cells expressed the highest levels of *NANOG* and *LRRC75A*. As others have reported (Chen et al., 2021), *PHC1* was highly expressed in hESC clusters R1-R3, and its expression was greatly reduced in differentiating cells. Downregulation of *PHC1* was compensated by increased *PHC2* expression, indicating a role for PHC2 in human neuronal differentiation (**Fig. 2E**). Focusing on the *FOX* family of TFs, *FOXD3*, which is required for pluripotency (Krishnakumar et al., 2016), and the recently reported pluripotency marker *FOXD3-AS1* (Haswell et al., 2021), were expressed in R1-R3 (**Fig. S3A**). Furthermore, a clear switch was observed from *FOXH1* and *FOXD3-AS1* expression in R1-R3, to the expression of the master regulator of brain development *FOXG1* (Beyer et al., 2013; Chiu et al., 2014) in all other clusters (**Fig. 2E** and **S3A)**.

### LSX forebrain induction cues evident at the end of Stage I

Under LSX induction, hESCs undergo morphogenetic events and form neural rosettes. These Day 7 cells mapped to a single cluster (R4, **Fig. 2D**), enriched in the rostral markers *SIX3* and *LIX1* (**Figs. 3, S2E** and **S3D**). In contrast, the expression of the preplacodal genes *EYA1* and *SIX1* (Ikeda et al., 2007; Schlosser, 2014) was low, including *SIX3-AS1*. Furthermore, expression of caudal markers *PAX5* and *GBX2* (Kirkeby et al., 2012; Maroof et al., 2013) was low throughout differentiation. This confirms the efficacy of LSX forebrain induction, enabling cell-fate commitment persistence.

Upon neural induction, the key neural TF PAX6 is upregulated and interacts with SOX2 (Zhang et al., 2019). R4 cells were *SOX2* positive showing high *PAX6* and *RAX* expression (**Fig. S2E)**, and express neuronal rosette markers, such as *DACH1, POU3F2, NR2F1* and *NR2F2* (Fedorova et al., 2019). A distinct switch from *CDH1* (Epithelial Cadherin) to *CDH2* (Neural Cadherin) expression was observed (**Fig. 2E**), and other developing forebrain specification and differentiation stage markers (such as *OTX2, HESX1, FOXG1, LIN28A* and *FABP7*) were detected.

### Self-patterning does not affect fate commitment

At Day 13, which marks the end of the self-patterning stage, 75% of the cells mapped to cluster R5 and most of them expressed *REST* (**Figs. 2E** and **S2E**). *SIX3, DLX5* and *BMP4* (**Figs. S2E** and **S3D**) were expressed in R6 cells that are enriched in *FGF8* and *HES1*. Moreover, R6 was enriched in *TAGLN*, which was also expressed in 44% of R5 cells but absent in the R5 cells co-expressing *NKX2*.*1, DCT* and *SOX6. CNTN1*, an active ligand of Notch (Hu et al., 2003) and potent inducer of neuronal migration (Lee et al., 2014), was expressed in R6. *CNTN1* was exclusively expressed in cells negative for *NTN1, DLL1, FABP7* and *POU3F2*. Comparing to results of 8 week human embryonic tissue (Kirkeby et al., 2012), no midbrain and hindbrain markers were detected at Day 13, confirming that the self-patterning phase does not affect fate commitment.

### Characterization of the Day 20 heterogenous population

Day 20 cells retained their identity and clustered in R7-13 (**Figs. S2E** and **3**). Some cells expressed high levels of *CDK1* (**Fig. S2E**), whereas other cells were still regulated by *REST* and expressed *DLX5* and *CDKN1C. CDKN1C*, which forms complexes with histone deacetylases to repress neuronal genes in non-neuronal cells (Laukoter et al., 2020) is inversely correlated with *REST* expression and enhanced in R11-13 (**Figs. 2E** and **3**). Interestingly, *ARX*, a regulator of cortical progenitor expansion by repression of *CDKN1C* (Colasante et al., 2015) was only expressed in R13 cells (**Fig. S2E)**. Neuronal differentiation correlated with *CDK6* upregulation and G1 shortening. *CDK6* is directly regulated by *GLI3* and expression of *GLI3* (Hasenpusch-Theil et al., 2018)(detectable at R4) dropped significantly in R12-13. *REST* is known to be downregulated during neurogenesis and in differentiating neurons and the pattern was recapitulated in this study (**Fig. 2E**).

### Neuronal maturation signatures

Day 20 cells were highly enriched for *MAP2*, and clusters R11-13 were enriched for *DCX*, which is a marker of migratory neurons. Genes expressed in proliferating neuroblasts associated with cortical migration control and developing rostral brain structural patterning, such as *EMX2* (Pang et al., 2008; Spalice et al., 2009; Verrotti et al., 2010), decreased in clusters R11-R12 and were undetectable in R13 (**Fig. S2E**). FGF8, an anterior-posterior patterning molecule, acting mainly via EMX2 repression (Hao et al., 2019), was expressed in R4 and R6 cells and in a few Day 20 cells, mainly in R8 and R10 clusters. Furthermore, *FGF17* (**Figs. 3** and **S2E**) and *FGF18* were mostly expressed at Day 20 R8 cluster. *HES6*-enriched cluster R11 (**Fig. S2E**) was composed of Day 13 and 20 cells, and most of the R11 *NEUROG1-*negative cells were Day 13 cells. Neural stem and progenitor marker *ZEB1*, which was downregulated upon neuronal differentiation to permit proper migration of immature neurons (Wang et al., 2019a), was expressed in almost all cells (**Figs. 3** and **S2E**). In addition, *FOXG1*-enriched R13 cells also express high levels of *DLX6-AS* and *DLX5* (**Figs. 3, S2E** and **S3B**).

Of note, the expression of *GNRH1* (Gonadotropin Releasing Hormone 1) was expressed in 30% of the R12-13 cells (9% of Day 20 cells) (**Figs. 3** and **S2E**). Of these cells, some expressed GABAergic or glutaminergic processing enzymes. As the mechanisms that contribute to the development of extrahypothalamic GnRH neurons are not fully described, such data are vital for studies of development, puberty and reproduction.

### Global expression profiles reveal neuronal differentiation and maturation signatures

To increase gene expression sensitivity, we performed bulk gene expression analysis with higher sequencing depth (**Fig. 4** and **S4**). Overall, we found 11,313 differentially expressed genes (DEGs) comparing cells from Day 0 to 20 (**Table S4**). More genes were differentially expressed during neural induction (Day 0 to 7), compared to the later stages, with self-patterning (Day 7 to 13) and maturation (Day 13 to 20) stages (**Table S4**). The most extensive transcriptional changes occurred between Day 0 and Stage I, with loss of pluripotency and gain of neuralization markers (**Figs. 4A, S4A and S4F**). We confirmed that bulk RNA-seq analysis for selected marker genes correlates well with ddPCR (**Fig. 4B** and **S4G**).

**Figure 4.**
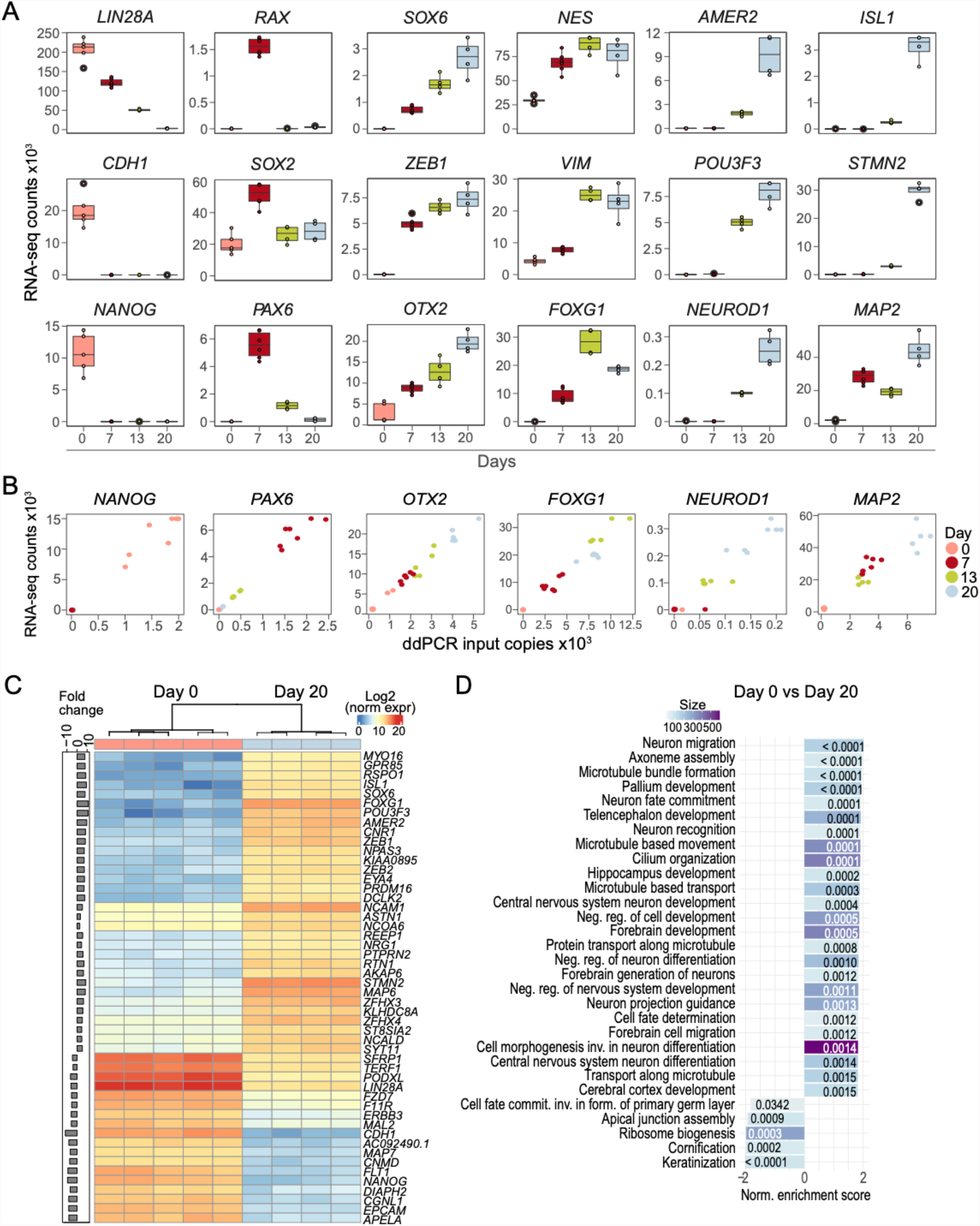
Correlation of RNA-seq to ddPCR and GO analyses results. A) Normalized gene expression counts for selected genes showing transcriptome expression patterns from loss of pluripotency towards neuronal maturation. B) Scatter plots of RNA-seq and ddPCR for marker genes *NANOG, PAX6, OTX2, FOXG1, NEUROD1* and *MAP2* at Days 0, 7, 13 and 20. C) Heatmap of top 50 differentially expressed genes between Days 0 and 20 replicates. Fold change is shown to the left. D) GSEA analysis of differentially expressed genes from Days 0 and 20.

The analysis shows the specific gene expression patterns as cells lose pluripotency and move towards neuronal maturation (**Fig. 4**). These may be steep decreases after neuronal induction, as seen for *LIN28A* and *CDH1*. For other genes expression peaks at Day 7 or Day 13, such as *RAX* and *FOXG1*, respectively. Expression increases gradually for genes such as *OTX2* and *SOX6*. Moreover, the expression of neuroectodermal patterning Wnt/β-Catenin negative regulator *AMER2* (Pfister et al., 2012), and neuronal differentiation marker *STMN2* (Wang et al., 2019b) increase at Day 13 and increase further at Day 20 (**Fig. 4A**). On Day 20 we also find genes correlated to specific neuronal types, such as *GRIA1, SLC17A6, GNRH1* and *GAP43* (**Fig. 4A** and **S4F**).

We next performed gene ontology (GO) analyses to identify shared biological processes (BP) among the DEGs during differentiation (**Fig. 4D** and **Table S5**). These analyses revealed enrichment of upregulated BPs related to pattern specification, neuronal maturation and migration from Day 0 to 20 (**Fig. 4D**). Stage-specific GO analyses revealed enrichment of BPs involved in neurogenesis and neuron development differentiation at stage I (Day 0 to 7), and BPs involving synaptic organization and signalling, and neurotransmitter regulation and secretion at the end of the maturation stage (Day 13 to 20; **Table S5**). The RNA-seq analysis is in line with single-cell analysis showing downregulation of pluripotency genes and upregulation of brain development genes.

### DNA methylation correlates with neuronal transcriptional programs during differentiation

DNAm in human cells is mainly restricted to CpG sites and essential for normal development (Smith and Meissner, 2013). As hESCs transition to differentiated neurons, dynamic DNAm changes regulate gene expression and the establishment of cell-type specificity (Stricker and Götz, 2018). To assess DNAm in the present protocol, we identified CpGs which are differentially methylated (DMCs) between Day 0, 7, 13 and 20 (**Figs. 5** and **S5**). As expected, comparing Day 0 and 20 reveals massive DNAm changes (n=210,049 DMCs, **Table S4**). Although we observe major changes in DNAm during the differentiation protocol (**Table S4**), the bulk DNAm levels and the distribution of unmethylated and methylated CpGs remains the same across all four timepoints (**Figs. 5A** and **S5B-C)**.

**Figure 5.**
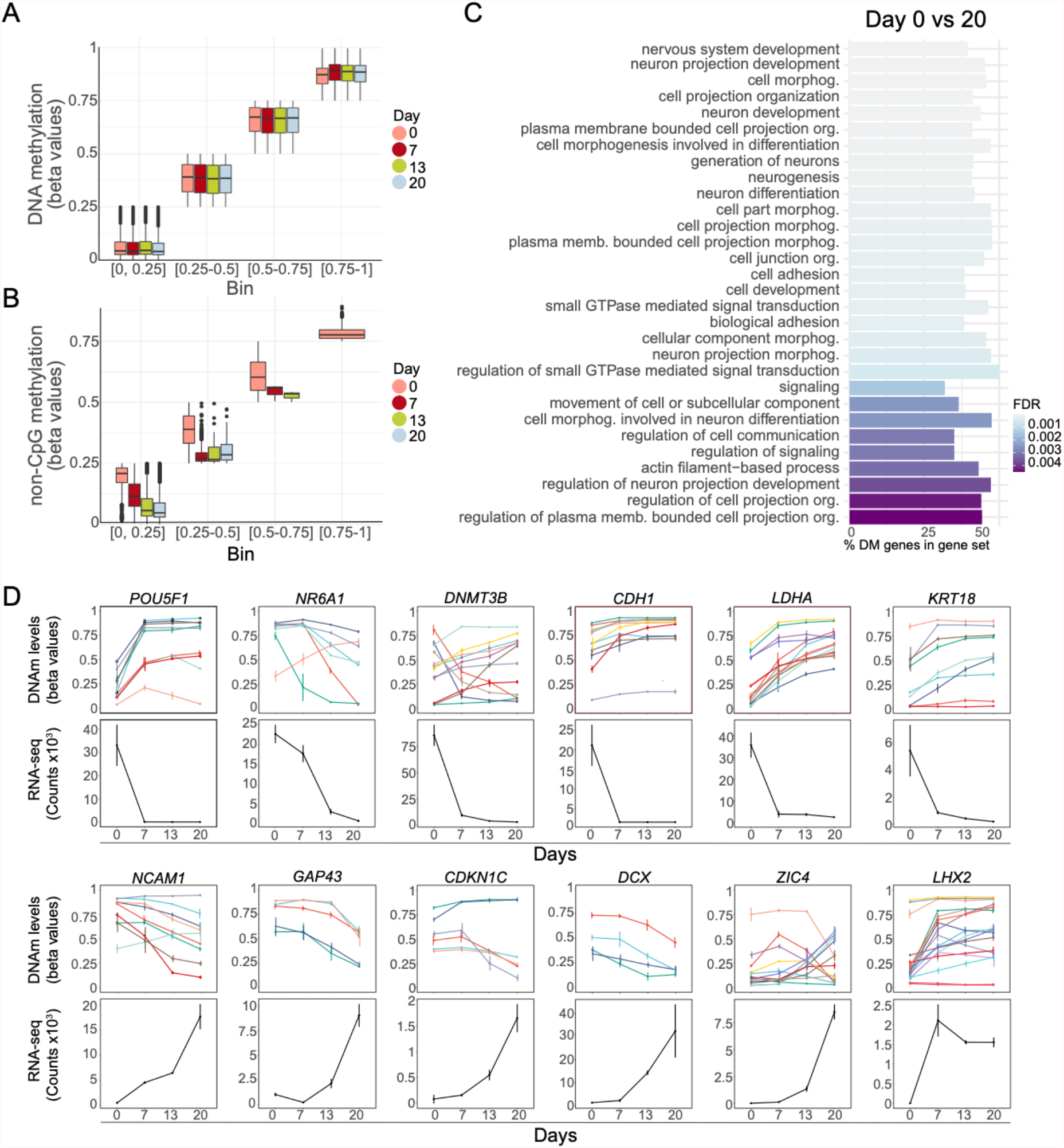
Specific DNAm changes during neuronal differentiation. A) Mean DNAm levels for each sample across all CpGs and non-CpGs (grouped in bins of 0.25) at Days 0, 7, 13, and 20. B) Mean DNAm levels for each sample across all non-CpGs (grouped in bins of 0.25) from Days 0, 7, 13 and 20. C) GOMETH analysis of top 30 BPs based on top 10% DMCs for Day 0 to 7. D) Significant CpGs of gene expression (derived from MORE) for Days 0, 7, 13 and 20. Top panels show DNAm mean +/- standard deviation whereas bottom panels show normalized RNA-seq counts for selected genes.

Deconvoluting these changes temporally, the highest number of DMCs was observed between Day 0 and 7 (n=161,600), with fewer changes in the self-patterning phase (Days 7-13, n=39,545) and during cell maturation (Days 13-20, n=47,676) (**Table S4**). Next, we used GOMETH analysis (Maksimovic et al., 2021) to explore shared biological functions among the DMCs. In line with the gene expression results, from Day 0 to 20 we observed enrichment of BPs involved in neural induction, neurogenesis, and brain development (**Fig. 5C**). This suggests that DNAm is modulating the neuronal transcriptional programs during the course of differentiation (**Fig. 5A**). Similarly, these analyses identified DMCs between Day 0 to 7 and Day 0 to 13, with BPs involved in cell adhesion and neuron projection morphology, which fits well with the stage cell transitions (**Fig. S5D** and **E**). One of the most significant GO terms is “neuron migration”, evidenced by expression of genes such as *DCX* and its partner *PAFAH1B1* (Nadarajah and Parnavelas, 2002) (**Fig. 5C**), both highly expressed in R11-13 (**Fig. 2** and **hESCNeuroDiffscRNA**).

To explore the correlation between DNAm and gene expression, we combined the DNAm and RNA-seq data sets based on CpG probe location and gene locus (**Figs. 5D, S57-G**, and **Table S4**). Of the Stage I gene annotated DMCs, 72% overlap with differentially expressed genes, inferring functional impact on gene expression. For genes with DMCs we generally observed a decrease upon transcriptional activation or an increase for genes becoming repressed during the course of differentiation. The expression levels of the majority of the differentially expressed genes between Day 0 and 20 are predicted to be associated with DNAm changes (8,011 of the 11,313 DEGs). The expression of markers of late trophectoderm (e.g., *KRT18)*, pluripotency maintenance (*POU5F1)*, suppression of pluripotency (*NR6A1)* (Wang et al., 2016), metabolic reprogramming (*LDHA)* (Zheng et al., 2016), or spatiotemporally regulated cortical TFs and cell cycle related genes (*LHX2, CDKN1C)* (Chou and Tole, 2019; Laukoter et al., 2020); and neuronal differentiation and maturation markers, (such as *DCX)*, may be regulated by one or more CpGs (**Fig. 5C, S5F, S5G** and **Table S6**).

Of note, the average non-CpG DNAm levels, and the distribution of unmethylated and methylated CpHs vary across time points. CpH DNAm is associated with transcriptional repression in the mouse genome (Xie et al., 2012) and non-CpG DNAm levels are enhanced at Day 0 cells and decline during differentiation (**Fig. 5B**).

### Chromatin accessibility analysis identifies regulation signatures during differentiation

To further assess the changes in the epigenetic landscape upon differentiation, we performed single-cell assay for transposase accessible chromatin sequencing (scATAC-seq). This analysis aimed at understanding chromatin-based gene regulation during neuronal differentiation from the loss of pluripotency at Day 0 to Day 20 (**Fig. 6** and **S6)**.

**Figure 6.**
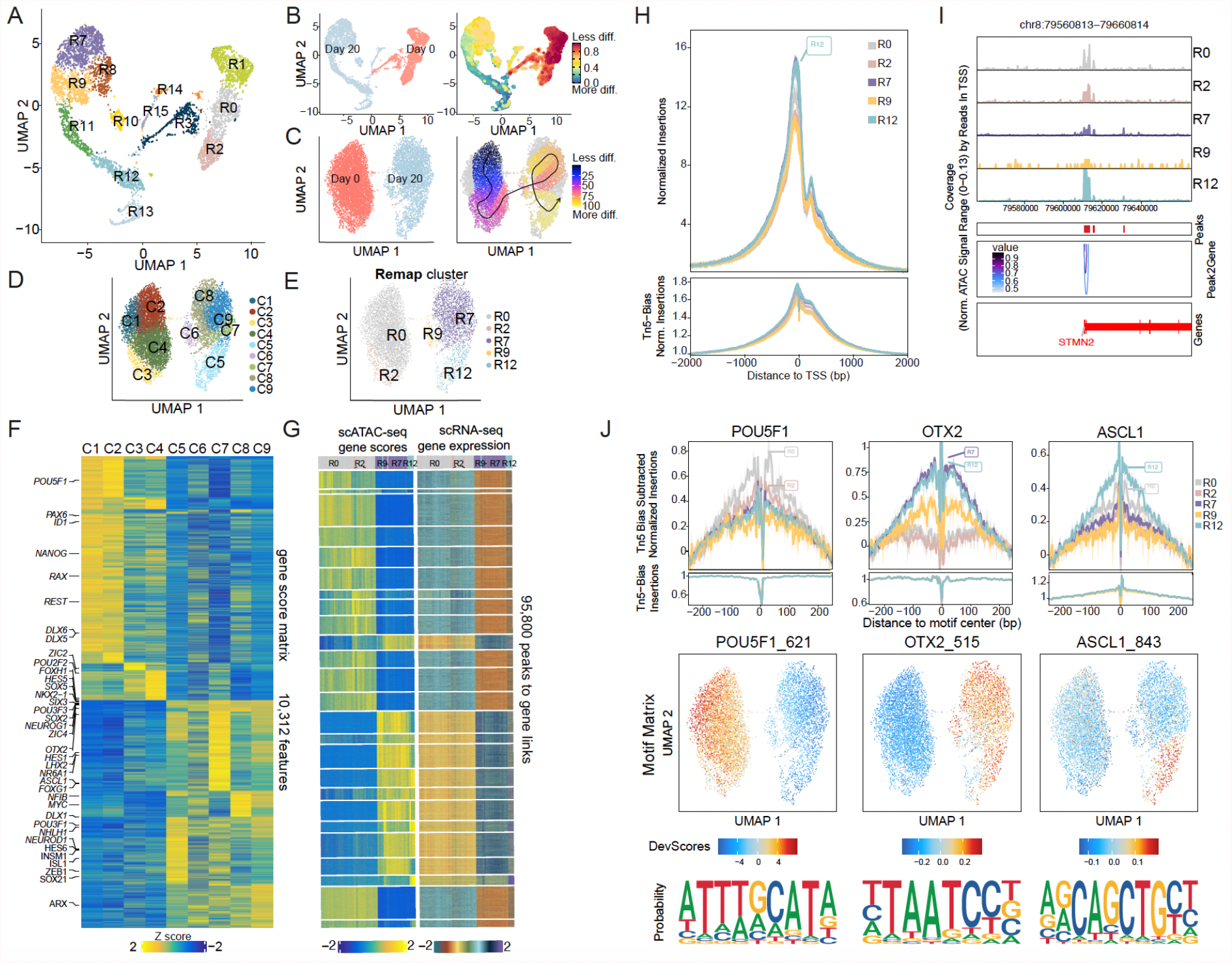
Integration of single-cell chromatin opening with scRNA-seq during neuronal differentiation. A) Multidimensional reduction UMAP plot of scRNA-seq corresponding to the timepoints used for scATAC-seq analysis. B) UMAP plot showing original identity of cells at scRNA-seq modality and corresponding differentiation trajectory. C) UMAP plot showing original identity of cells at scATAC-seq modality and corresponding supervised pseudo time trajectories. D) UMAP plot showing clusters at scATAC-seq modality. E) Remap UMAP plot of renamed clusters following constrained alignment of cell populations after integration of scATAC-seq and scRNA-seq. F) Top selected marker genes from scRNA-seq data shown on a heatmap plot computed on Gene Score Matrix. G) Peak to gene linkage heatmap for scRNA-seq clusters and corresponding gene scores for integrated scATAC-seq clusters. H) Chromatin openness of integrative cluster R0, R2, R7, R9 and R12 over all TSS. I) Tracks shown on peak browser for selected gene *STMN2* on integrated cell clusters. Bottom panel shows co-accessibility interactions around TSS. J) Motif footprinting for selected transcription factors POU5F1, ASCL1 and OTX2 demonstrating preferential opening in different cell clusters. The middle panel shows the corresponding motif deviation scores of ArchR identified TFs POU5F1, ASCL1 and OTX2. The scores are calculated for each TF motif observed in an accessible region and in each cell for the deviation from expected average accessibility across all the cells. The representative sequence logos identified in accessible regions across the dataset are shown below.

### Reanalyzing scRNA-seq datasets for integration with scATAC-seq data

To integrate scATAC-seq and scRNA-seq, the scRNA-seq datasets for Day 0 and Day 20 were reanalyzed. 1910 Day 0, and 3033 Day 20 cells were projected in 14 clusters (**Fig. 6A**) and in accordance with the maturation trajectory seen in the corresponding CytoTRACE plot (**Fig. 6B**). The scRNA-seq clusters were numbered and annotated similarly to old clusters (**Fig. 6A, S6A** and **B**). Day 0 cells resolved into five clusters (R0-3 and R14) whereas Day 20 clusters were numbered as R7-15, cohering to the initial four-timepoint analysis.

### Chromatin accessibility changes globally during differentiation

The analysis of 4,901 Day 0 nuclei and 2,847 Day 20 nuclei and the scATAC-seq data showed a good distribution of fragment sizes, fragment numbers, and TSS enrichment (**Fig. S6C, D, E and F**). The supervised pseudotime trajectory analysis, which predicts paths for gene regulatory changes in cells during differentiation, showed a similar profile to the gene expression CYTOTRACE analysis (**Fig. 6B** and **C**). We mapped four chromatin accessibility clusters at Day 0 (C1-4) and five at Day 20 (C5-9; **Fig. 6D**) and observed differential chromatin opening in these cell clusters for many loci, including *POU5F1, REST, GAD2*, and *DCT* (**Figs. 2, S2, S3** and **S6G**). We next generated a gene score matrix heatmap incorporating regulatory elements, representing a score of chromatin opening of 200 kb gene regions. The heatmap shows a selection of marker genes based on their relevance to pluripotency and brain development and the previously described scRNA-seq cluster annotation (**Fig. 6F)**. Higher gene scores for genes that are known for their role in the regulation of pluripotency, such as *POU5F1, NANOG, ID1*, and known enhancer specific binding factors in development, such as *ZIC2* (Hong et al., 2011) were found in Day 0 clusters (**Fig. 6F, 2 and S6B**). *SOX2* is regulated by several enhancers and interacts with multiple but distinct groups of transcription factors, including POU3 class partners (Iida et al., 2020; Mistri et al., 2015; Tang et al., 2015; Zhu et al., 2014). Chromatin accessibility for *SOX2, POU3F1*/*BRN1* and *POU3F3*/*BRN3* increased with differentiation (**Fig. 6F**). Moreover, genes expressed at the differentiation endpoint clusters, such as *ASCL1* and *SOX21* which are implicated in neurogenesis (**Fig. 2 and S3**), *NFIB* which is crucial in neural progenitor cell renewal (Piper et al., 2014), and *OTX2* which is associated with early neuronal development regulation, showed a more open chromatin structure in neuronal clusters C5-C9. The gene score of *NEUROD1* is highest in the trajectory end-point cluster C5 (**Fig. 6F)**.

### Correlation of chromatin regulatory dynamics and gene expression

To better understand the regulatory interactions with gene expression we performed integrated analysis of scATAC-seq with scRNA-seq using ArchR (Granja et al., 2021). Following constrained alignment of cell populations after integration of scATAC-seq and scRNA-seq, the integrated clusters were renamed to correspond to the previously annotated scRNA-seq clusters **(Figs. 6E** and **S6B**). Pluripotency clusters C1-C4 remapped to scRNA-seq Day 0 clusters R0 and R2, whereas clusters C7-C9 mapped to cluster R7, correlating chromatin openness and gene expression in single cells for markers such as *REST, HES1*, and *CDK1* (**Figs. 6B, D, E, S6B** and **S6I**). Cluster C6 mapped to R9, which was marked by expression of *REST, HES5* and *ASCL1*, and C5 remapped to one of the endpoint clusters, R12, having high *NTRK1* expression (**Fig. S6I**). We assessed scATAC-seq peaks across *TGDF1, CDH1, CDH2, STMN2*, and *DCX* loci across the integrated clusters and found cluster specific chromatin opening (**Fig. 6I** and **S6J**). Moreover, the peak-to-gene co-accessibility arcs show gene expression linked to chromatin opening during differentiation of putative distal regulatory elements at *TGDF1* and *CDH1*.

To explore the integration of chromatin accessible regions and gene expression, we mapped 95,800 peak-to-gene links and observed a clear correlation of chromatin regulatory dynamics in the different integrated clusters (**Fig. 6G**). Moreover, chromatin accessibility was clearly enriched at transcriptional start sites (TSS) in every cluster (**Fig. 6H**). However, chromatin accessibility peak annotation analysis revealed varying enrichment across integrated clusters at promoters, intronic, exonic and distal regions (**Fig. S6H**). The number of peaks at promoters were very similar across the five clusters, with fewer peaks in cluster R9. Interestingly, more peaks were detected at intronic and distal regions in clusters R7 and R12 than in the other clusters. These results agree with previous studies showing that neuronal gene activation depends on multiple regulatory regions, many of which are located far from the gene locus itself.

We next aimed to explore the dynamics of lineage-defining factors at pluripotency and differentiation endpoint. Using ArchR, we identified specific TF motifs across differentiation (**Fig. 6J**). Motif footprinting for *POU5F1* underlies a regulatory function in accessible chromatin in pluripotent clusters R0 and R2, whereas motifs for *ASCL1* and *OTX2* footprints were more enriched in differentiated clusters R7 and R12. We further mapped the enrichment of *POU5F1, DLX6, ASCL1* and *OTX2* motifs in open chromatin in individual cells (**Figs. 6J** and **S6K)**. These examples illustrate how lineage-defining TFs dynamically regulate gene expression programs during neuronal differentiation.

### Exploration of single-cell data using interactive webtools

Large dataset analyses, such as single-cell sequence analysis, generally require bioinformatics expertise for interpretation. We have made our scRNA- and scATAC-seq data accessible to a broader audience by providing open access web-interfaces based on open-source tools, abiding by the Findability, Accessibility, Interoperability, and Reusability (FAIR) principles (Ouyang et al., 2021; Sharma et al., 2021). The users can explore scRNA-seq data in **<u>hESCNeuroDiffscRNA</u>** and plot high resolution figures of their genes of interest under seven different tabs (**Fig S7A-H**). This includes exploration of 1) Gene expression UMAPS as illustrated for *POU5F1* and *NTRK1*; 2) gene co-expression analysis, here shown for *PHC1/PHC2* and *NEUROG1/NTRK1*; 3) different gene and cluster expression configurations, such as heatmaps, violin-, box-, proportion- and bubble plots. The platform also allows for correlation with other published gene expression datasets (**Fig. S7H**).

To illustrate the utility of the web interface, we focus on *ZIC2* and *ZIC4*, and their expression and regulation during neuronal differentiation (**Fig. 7**). ZIC proteins are known for their role in proliferation and differentiation of neural progenitors, neurulation, neural tube formation, and neural plate closure (Al-Naama et al., 2020; Aruga and Millen, 2018). Global expression analysis shows that *ZIC2* is present at Day 0 and peaks at Day 7, whereas expression of *ZIC4*, mostly undetectable at early timepoints, appears at Day 13 and peaks at Day 20 (**Fig. 7A**). DNAm levels at the CpGs in the *ZIC2* locus were stable across differentiation. In contrast, DNAm of 13 CpGs in the *ZIC4* locus were positively or negatively correlated with gene expression across differentiation (**Fig. 7B**). Differential expression of *ZIC2* and *ZIC4* across the individual cells at Day 0, 7, 13 and 20 in UMAPs (**Fig. 7C**) can be compared and correlated with selected TFs, shown here to be important for regulation in the neuronal differentiation protocol (**Fig. 7C**).

**Figure 7.**
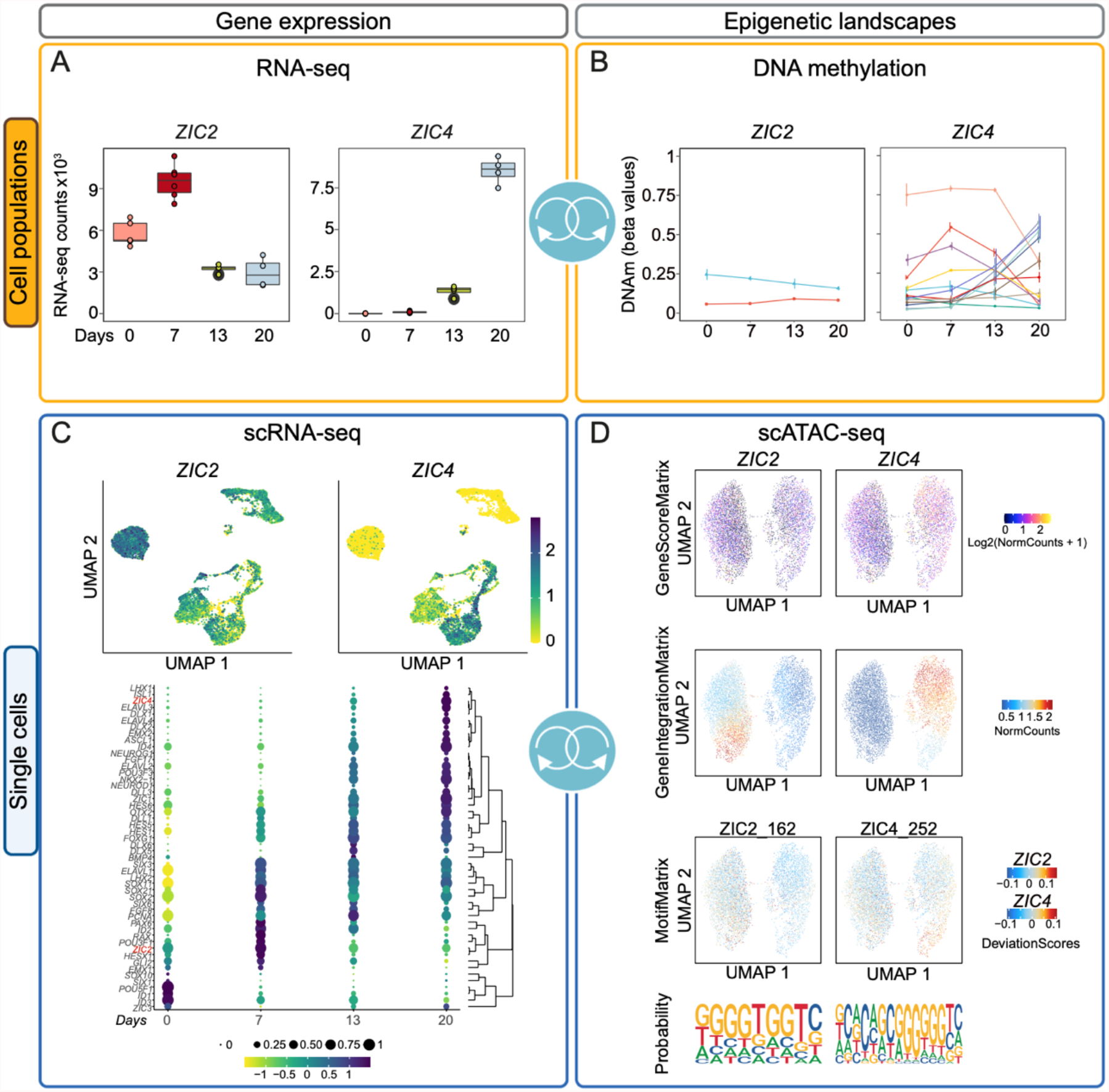
An example of the possibilities and potential applications of the 4D data showcasing the *ZIC2* and *ZIC4* genes. A) Normalized global gene expression counts for *ZIC2* and *ZIC4* from Days 0, 7, 13 and 20. B) Significant CpGs in gene loci *ZIC2* and *ZIC4* (derived from MORE) for Days 0, 7, 13 and 20. DNAm is represented as mean +/- standard deviation. C) Representative UMAPs showing cluster specific and differentiation-driven gene expression across all four timepoints for *ZIC2* and *ZIC4*. A bubble plot representing gene expression and hierarchical clustering of TFs highly relevant to neuronal differentiation across Days 0, 7, 13 and 20. D). The upper UMAPS represent inferred gene scores of the openness of the *ZIC2* and *ZIC4* gene loci. In middle UMAPs the gene integration shows correlation of gene expression with chromatin opening of the *ZIC2* and *ZIC4* gene loci. The lower UMAPs represent motif footprinting demonstrating preferential opening in different cell clusters for ZIC2 and ZIC4 with the representative sequence logos identified in accessible regions in the dataset below.

The scATAC-seq data can be explored in “**hESC Neuro Differentiation scATAC seq**” (**hESCNeuroDiffscATAC)** (**Fig. S7I-N**). Users can visualize chromatin accessibility, motif enrichment or integration of scATAC-seq with scRNA-seq in UMAPs. Furthermore, this webtool enables investigation of the gene score and motif matrix, also showing the representative sequence logo calculated from open regions. (Examples are given for the *L1TD1, HES5* and *ZEB1* genes at **Fig. S7J**). Chromatin opening can be explored across the genome for different clusters, as shown for *SOX2* and *POU5F1* or in different heatmaps (**Fig. S7K-N**). Heatmap views could either be pseudotime trajectories or peak-to-gene linkage, which define linked chromatin opening peaks with promoters of expressed genes (Granja et al., 2021) and may deduce enhancer promoter interactions (Baek and Lee, 2020). In the case analysis of *ZIC2* and *ZIC4*, gene score and gene integration analysis showed the genes were active in different cells (**Fig. 7D**) and the ZIC2 and ZIC4 footprints and representative sequence logos were identified. This 4D example analysis highlights the epigenetic regulation and gene expression of these genes in neuronal differentiation.

## Discussion

Here we present comprehensive multi-omics analyses to characterize a novel neuronal differentiation protocol from pluripotent ESCs towards a ventrally committed, telencephalic population of progenitors, mature and immature neurons. We assessed stage-to-stage transition using ddPCR and immunofluorescence imaging, and used scRNA-seq and bulk RNA-seq to validate cell populations over time. scATAC-seq and DNAm analysis further characterized the epigenetic and gene regulatory landscapes.

The deconvolution of early human neuronal differentiation at the level of molecular regulation provides insight to an otherwise inaccessible developmental window. Animal models are valuable, but evidence shows that the human neocortex develops under the effect of additional mechanism (Massimo and Long, 2021; Pinson and Huttner, 2021; Xing et al., 2021). Thus, neuronal differentiation studies from PSCs provide an alternative method to characterize developmental transcriptome trajectories and the roles of specific genes in human brain formation and patterning.

To specify the patterning and maturation identities of the cells, we followed up the trajectories of major TFs (O’Leary and Sahara, 2008). Ventral telencephalic markers, such as EMX2 and ASCL1, were already expressed at the end of Stage II, while dorsal telencephalic markers such as EMX1 and NEUROG2 were absent. Absence of expression of HOXB2, PAX7 and GBX2 at any timepoint, confirmed that the self-patterning phase after neural induction, had no effect on lineage commitment and no cells differentiated to hindbrain, midbrain or thalamic lineages.

GO analyses revealed stage dependent enrichment of biological processes correlated to neurogenesis, pattern specification, signalling and neurotransmitter regulation, migration, synaptic organization and neuronal maturation. The DNAm analyses showed alternating, stage-dependent changes for various patterning genes and TFs important to neurogenesis. In most developmentally regulated genes involved in neuronal lineage commitment, DNAm levels decreased upon transcriptional activation and increased for genes becoming repressed during the time of differentiation. We also observed that sometimes downregulation of gene expression in the self-patterning stage might intersect the upregulation of gene expression seen both in stage I of neural induction of hESCs and during maturation at stage III. This could be due to the combined effect of non-CpG and CpG DNAm implicated in the regulation of RNA splicing in ESCs and neurons, respectively (Ball et al., 2009; Laurent et al., 2010). Non-CpG DNAm accumulates in neurons during synaptogenesis and synaptic pruning (Lister et al., 2013), but CpH DNAm is associated with transcriptional repression (Xie et al., 2012). Whether and how CpH DNAm plays a role in self patterning following the LSX induction is not known, and future studies are needed to explore this. Furthermore, the *in vitro* model for DNAm changes presented here, is advantageous for neuropharmacological studies. Whether these changes can be translated to distinct early developmental events, cannot be ascertained. The direction of causality of epigenetic regulation for early brain development, can however further correlate *in vitro* models to sets of open cis-regulatory elements and the regulation of TF-centred networks. The identification of common DNAm modification sites and chromatin openness regions may present candidate loci for future studies of early human development and may advance translational studies of the impact of drugs used early in human pregnancy.

The effect of loss of pluripotency towards neuralization, irrespective of the intermediate timepoints, was investigated through an integrative analysis of the scATAC- and scRNA-seq. We used ArchR (Granja et al., 2021) for this analysis as this pipeline was flexible for small ATAC-seq and scRNA-seq datasets. The juxtaposition of the transcriptome to the regulatory elements in ESC and differentiating Day 20 cells can infer gene regulatory network information. We identified linked sets of genes unique to each state, that may comprehensively profile individual cells. Such approaches are highly informative. They can infer epigenome causality not restricted to these analyses, but also to studies of the effect of drugs in early brain physiology and development.

### Strengths of study

Although numerous studies have used the LSX cocktail for neural induction, to our knowledge this is the first study that has shared all scRNA data in such transparent and interactive format. Thus, a strength of this study is the presentation of our single-cell data in two visualization tools, ShinyCell and inhouse developed ShinyArchR.UiO (Ouyang et al., 2021; Sharma et al., 2021), that are openly available for users. These tools allow the users to explore candidate genes and utilize a comprehensive set of functionalities, beyond the fate specification analysis presented here. Furthermore, these tools enable insight into the molecular and structural partners of stage-specific markers and time-stamped TFs, their transcriptional regulation and cell cluster identities. Programming scripts for data analysis are made available and can be easily customized for further studies and the incorporation of other data. Although the protocol does not generate terminally differentiated neurons of a specific subtype, there are numerous advantages. The protocol is cost-effective at the level of culture coating reagents, vessel size and timed passaging. Moreover, the advantage of 2D culture, defined cell numbers at all passaging steps, reduces the human errors in reproducibility compared to protocols based on confluency evaluation and cumbersome 3D culture setup. As the protocol was designed for neuropharmacological studies, the daily media changes further diminish the effect of short drug half lives in such studies.

### Limitations of study

This study has several limitations at the level of cell characterization. The membrane electrochemical and electrophysiological maturation properties were not evaluated. Moreover, we did not assess neuropeptide diversity, secretion of neurotransmitters or migration. Protein quantification or intracellular localization of markers and trajectories were also beyond the scope of a multi-omics characterization. A limitation of scATAC-seq is the genome-per-cell coverage and open chromatin regions relevant for the individual cell or cell populations may have been missed.

In conclusion, in this study we describe the generation of a novel neuronal differentiation protocol where we used the unparalleled power of multi-omics to understand early events of anterior neuroectodermal fate specification. We assessed the functional regulation of transcription factors and developmentally regulated genes, from loss of pluripotency towards neuronal differentiation. Integration of scATAC-seq and scRNA-seq provide invaluable insight on the complexity of fate decisions and enabling other researchers to finetune future studies. Finally, the reader has access to the single-cell sequencing data in two searchable, user-friendly webtools to visualize intra- and inter-timepoint and cell cluster regulation, interactively.

## Acknowledgements

We thank Marit Ledsaak and Naima Azouzi for technical assistance. The majority of informatic analysis was performed at saga super computing resources (Project NN9632K) provided by UNINETT Sigma2—the National Infrastructure for High Performance Computing and Data Storage in Norway. The sequencing service was provided by the Norwegian Sequencing Centre (www.sequencing.uio.no), a national technology platform hosted by Oslo University Hospital and the University of Oslo and supported by the “Functional Genomics” and “Infrastructure” programs of the Research Council of Norway and the Southeastern Regional Health Authorities. PharmaTox Strategic Research Initiative was supported by Faculty of Mathematics and Natural Sciences, University of Oslo. We acknowledge funding from Research Council of Norway 262484 (R.E.) and 241117 (R.L.); the Swedish Research Council 2019-01157 (A.S.) and grants from the Swedish Brain FO2019-0087 (A.S.) and the Freemasons Children’s House of Stockholm (A.S.). The graphical abstract was generated in Biorender.com.

## Author Contributions

Conceptualization, A.S., K.G., R.L., R.E.; Methodology, A.S., M.S., R.L., S.M., K.G.,R.E., A.Sh., M.L., M.F.; Writing – Original Draft, A.S., M.S., A.Sh., M.L., K.G., R.L., R.E.; Writing – Review & Editing, A.S., M.S. R.L., K.G., R.E., A.Sh., M.L., M.F., G.A.; Software, Formal Analysis, and Visualization, A.Sh., M.S., K.G., P.M.B., R.L., R.E.; Investigation and Validation, A.S., M.F., M.S., M.L., S.M., R.E.; Funding, R.L., A.S., and R.E., Supervision, R.L., K.G., A.S., and R.E. Resources G.A., R.L. and R.E.

## Star methods

### KEY RESOURCES TABLE

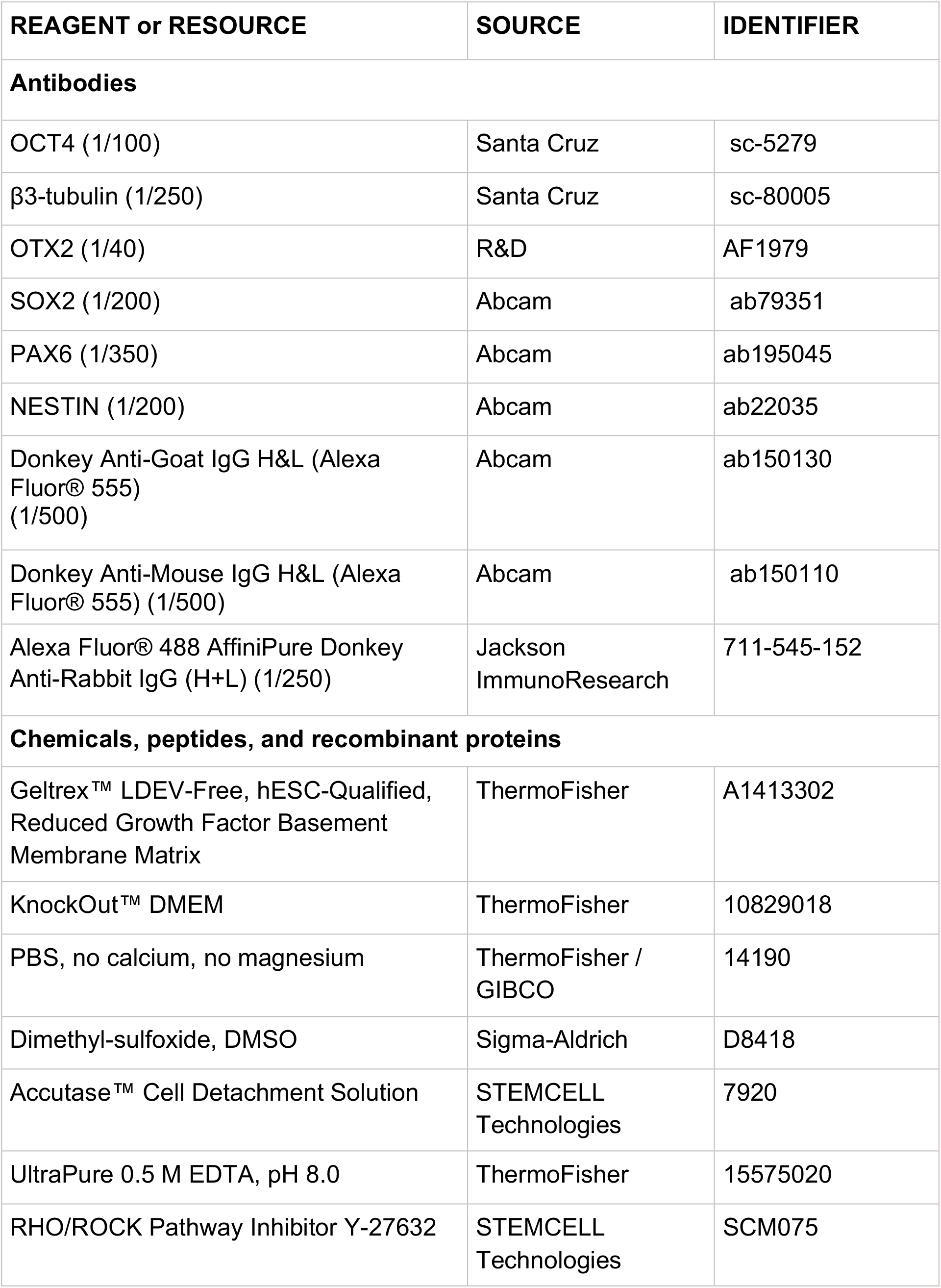
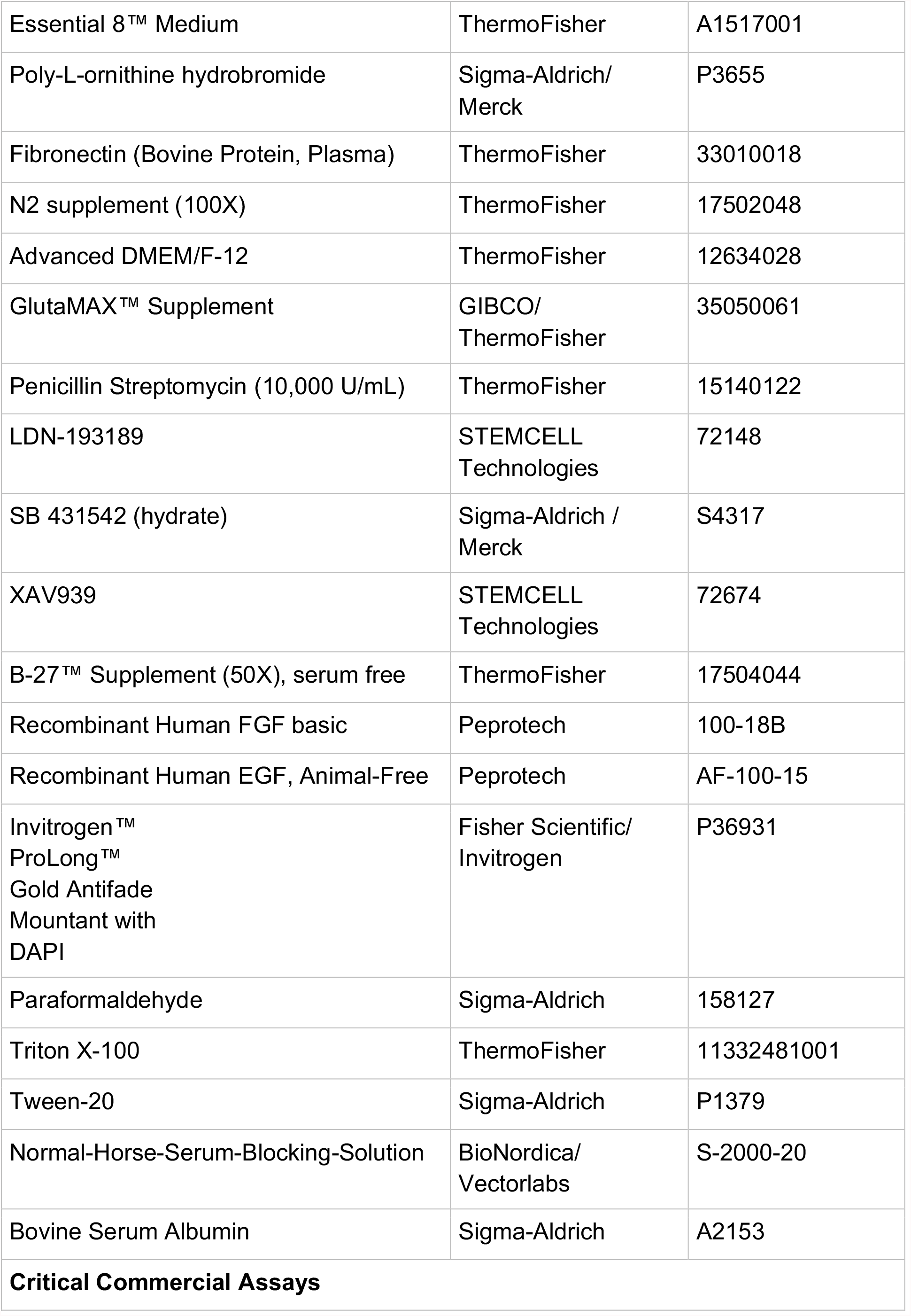
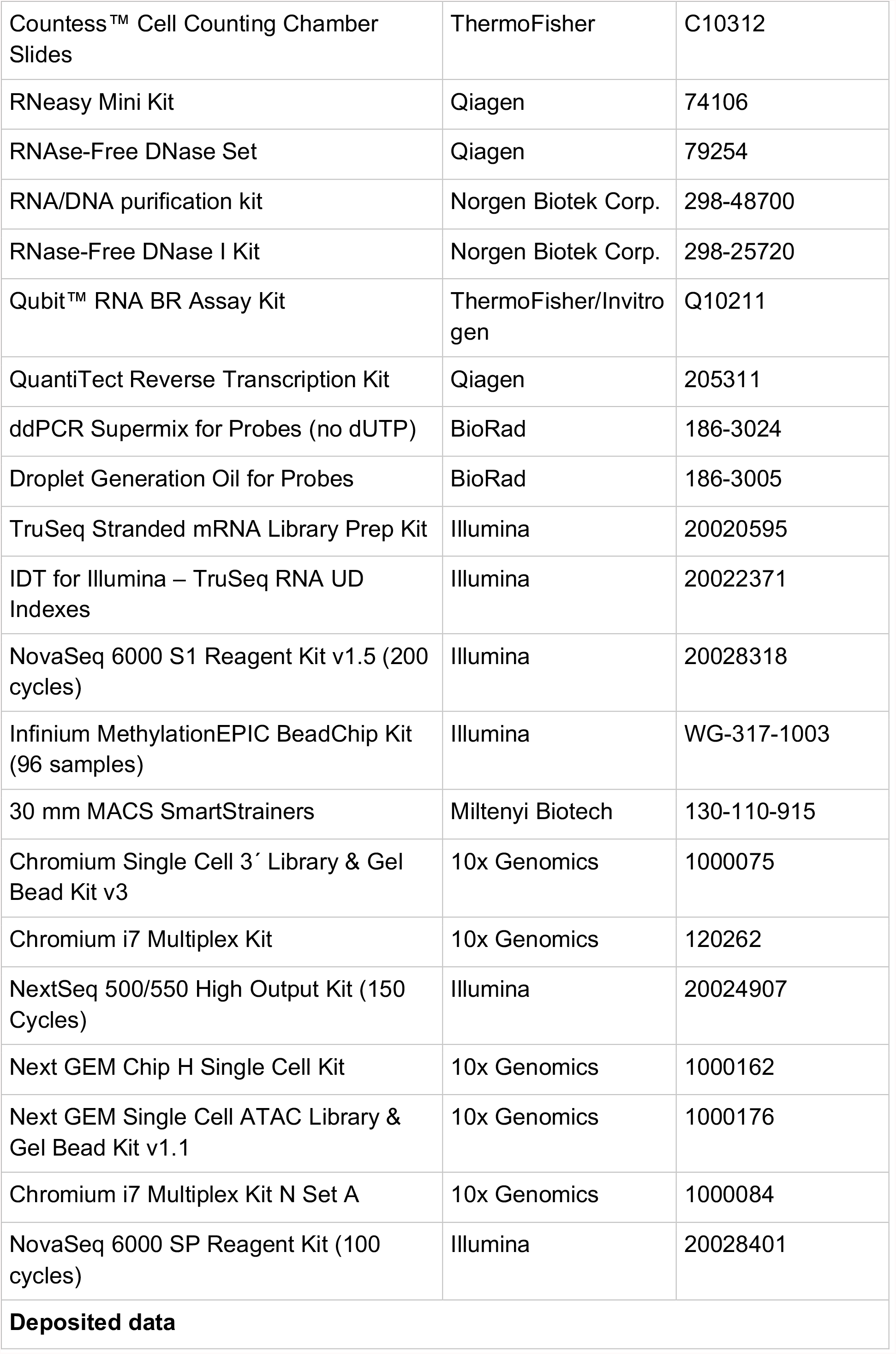
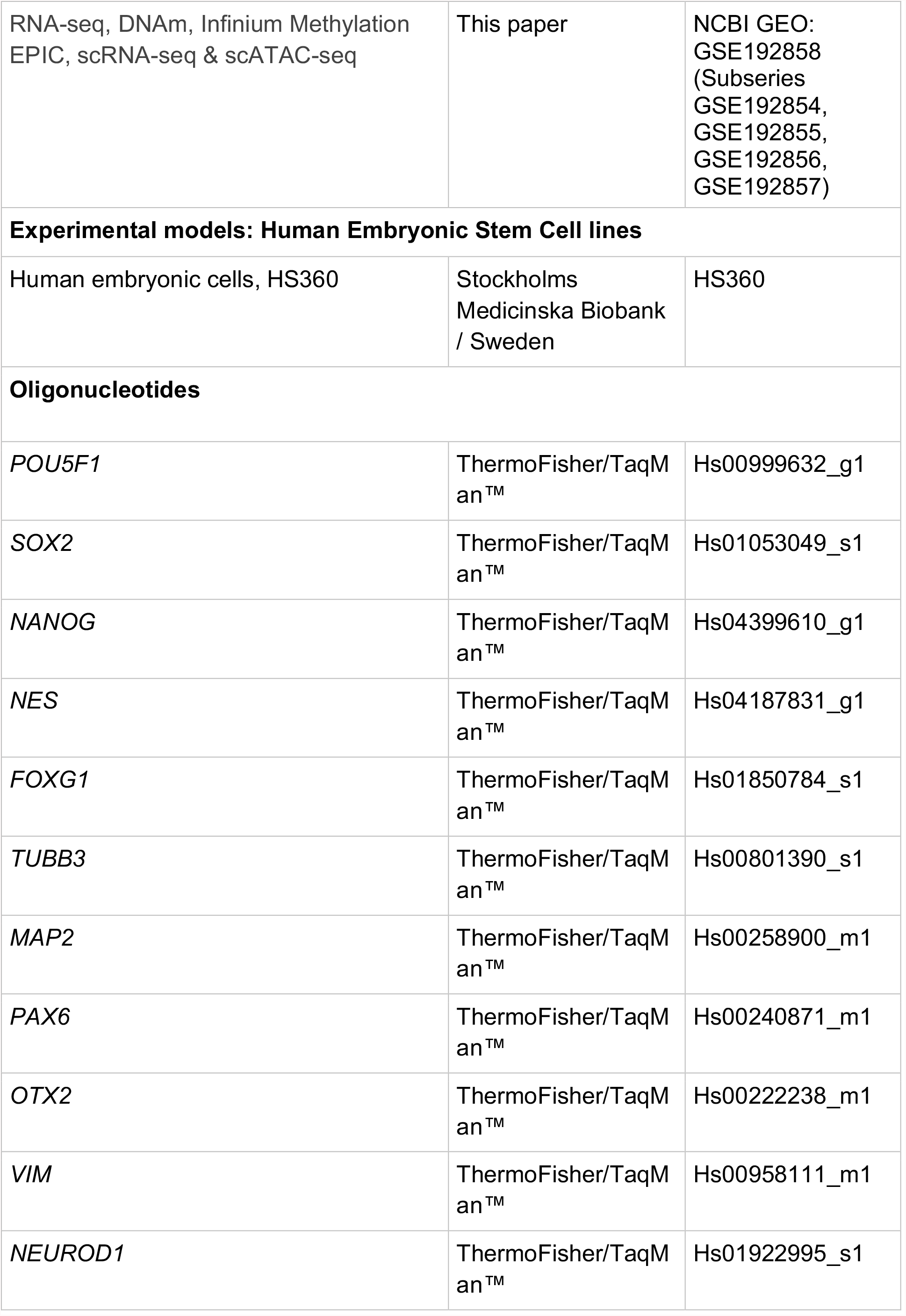
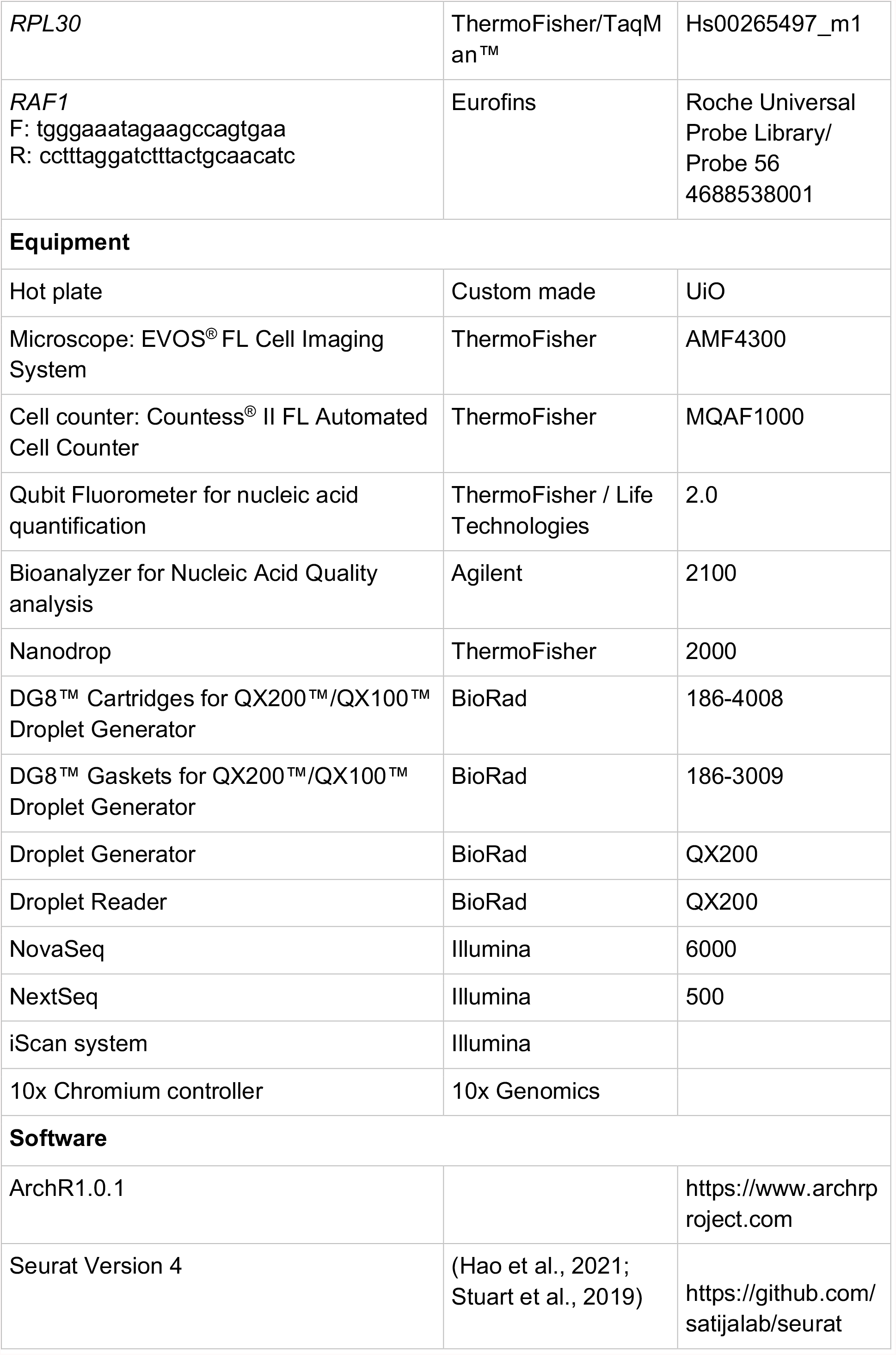
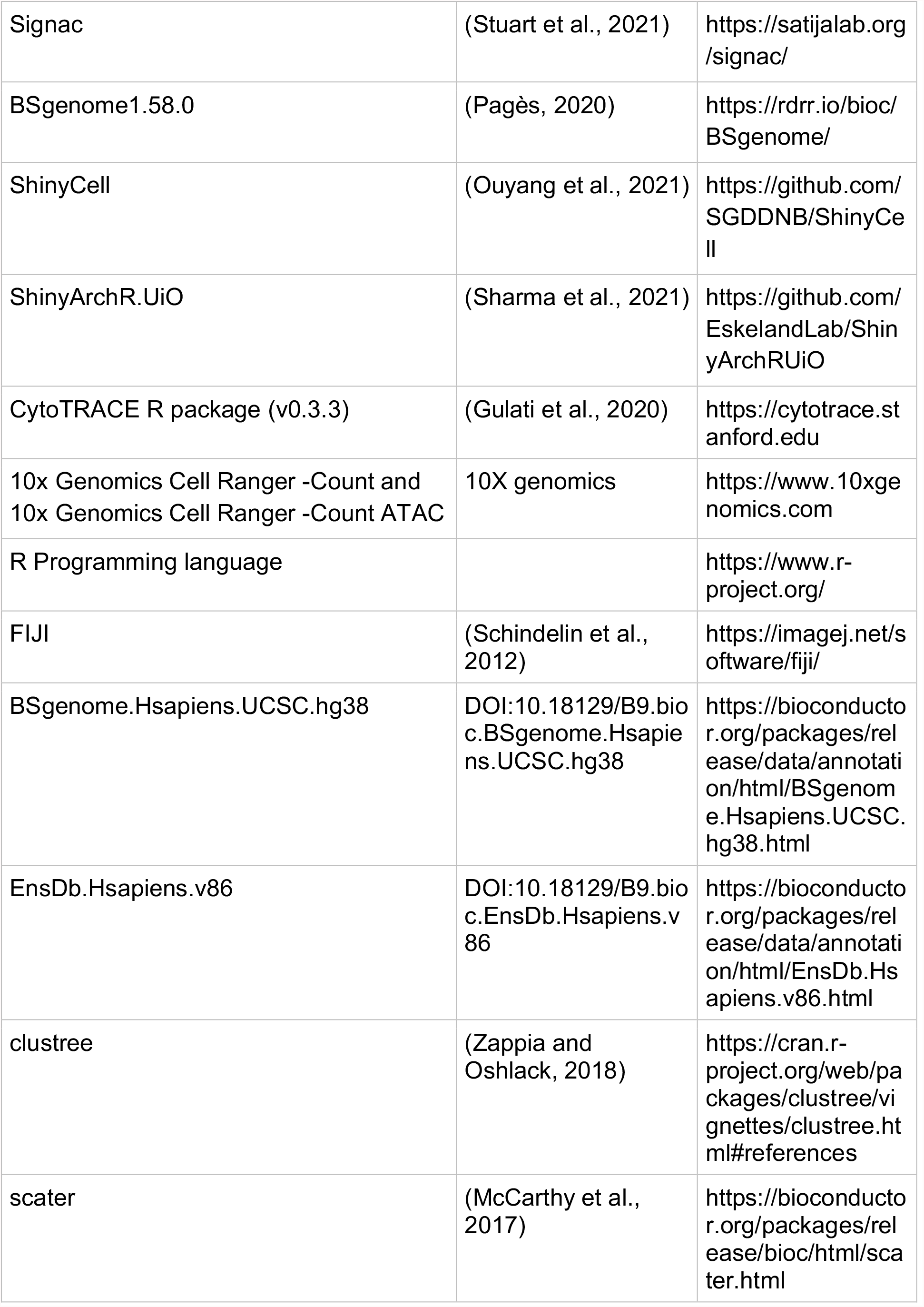
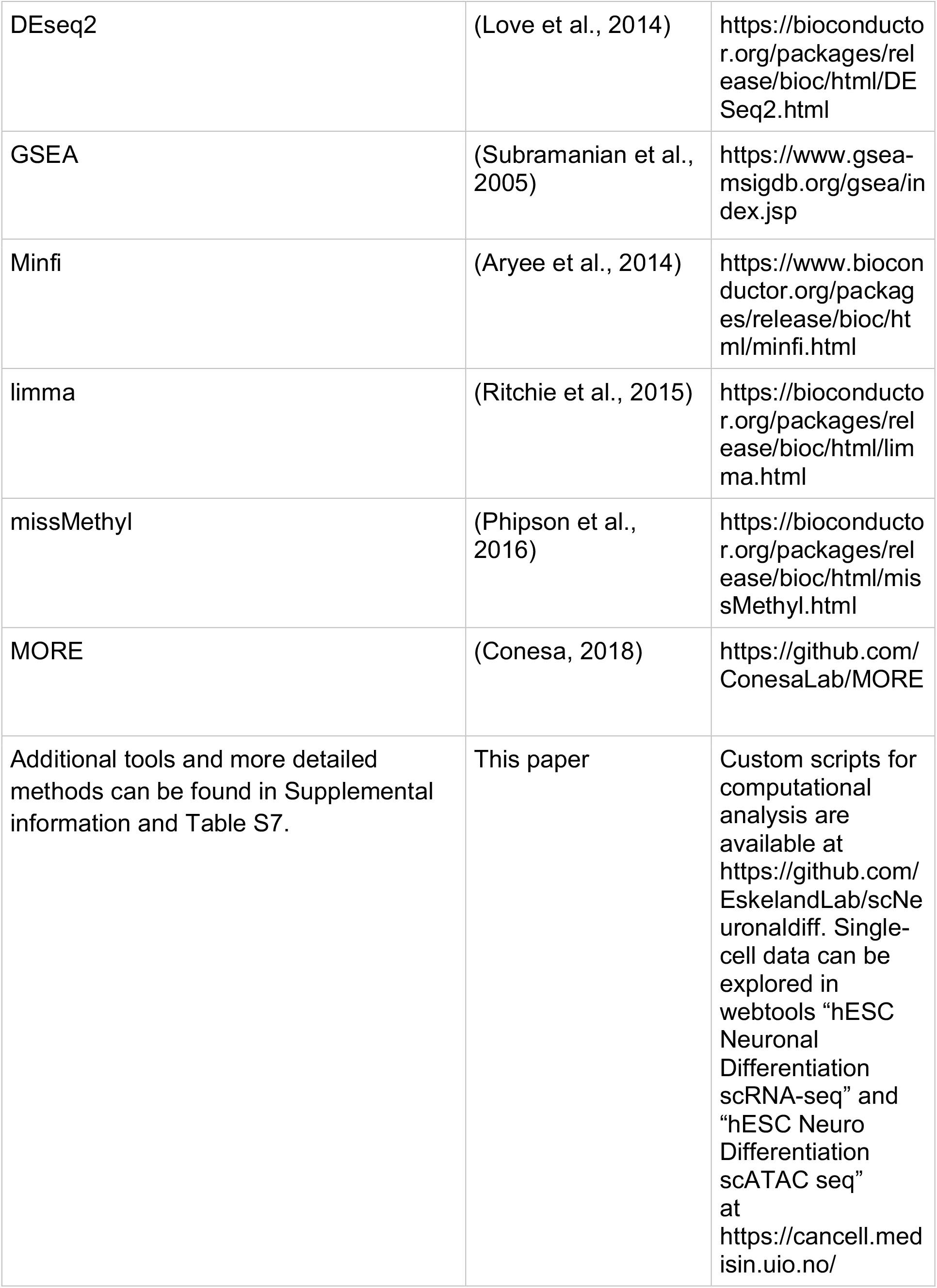

### RESOURCE AVAILABILITY

#### Lead contact

Further information and requests for resources and reagents should be directed to and will be fulfilled by the lead contact Ragnhild Eskeland.

### EXPERIMENTAL MODEL AND SUBJECT DETAILS

#### Human embryonic stem cell (hESC) culture and maintenance and neuronal differentiation protocol

The full description of the differentiation protocol is described in our protocol manuscript (bioRxiv https://doi.org/10.1101/2022.01.26.477818).

#### Immunofluorescence analysis

In brief, cells grown on 13mm glass coverslips, were washed once and fixed in 4% paraformaldehyde for 15 min at room temperature (RT). After 3 washes, the cells were permeabilized with 0.3% Triton X-100 (ThermoFisher) in blocking buffer containing 2% BSA (Sigma-Aldrich) and 0.01% Tween in 1×PBS for 30 min at RT, washed 3 times, and blocked with 10% horse serum for 30 min. Primary antibodies were diluted (as in KRT) in 1×PBS containing 0.03% Triton X-100, and coverslips were incubated overnight at 4°C. Next, coverslips were equilibrated at RT for 2 hours and washed 3 times. The secondary antibodies were diluted (see KRT) in 0.01% Tween-20 (Sigma-Aldrich) and 0.1% horse serum (BioNordica) in 1×PBS, and coverslips were incubated for one hour at RT. The coverslips were washed 3 times and mounted on microscope slides using the ProLong™ Gold Antifade Mountant containing DAPI (Fisher Scientific) to counterstain cell nuclei. Washing steps lasted 15 minutes and used 1×PBS. Images were obtained with a DeltaVision high resolution widefield microscope (GE Life Sciences, USA) using the Resolve 3D software and 100X 1.45NA oil objective and processed using the open-source software Fiji (Schindelin et al., 2012).

#### DNA/RNA isolation

Genomic DNA and total RNA were isolated by direct lysis in the culture well followed by column-based isolation using RNA/DNA purification kit (Norgen Biotek). The RNase-Free DNase I Kit (Norgen Biotek) was applied for on-column removal of genomic DNA contamination from RNA isolates. Three RNA isolates were processed using RNeasy Mini Kit (Qiagen) followed by DNase-treatment using RNAse-Free DNase Set (Qiagen). All isolations were done according to the manufacturer’s instructions. Nucleic acid quantification was performed using Qubit (ThermoFisher Scientific), purity was measured using Nanodrop 2000 (ThermoFisher Scientific), while RNA and DNA integrity was assessed using 2100 Bioanalyzer (Agilent Technologies) and 4200 TapeStation (Agilent Technologies), respectively.

#### Droplet Digital RT-PCR and RNA expression analysis

Reverse transcription of total RNA was performed using QuantiTect Reverse Transcription Kit (Qiagen). Subsequent ddPCR reactions were set up using ddPCR Supermix for Probes (No dUTP) (BioRad) and Taqman assays (ThermoFisher) or Universal Probes (Roche) in combination with target primers (Eurofins) as outlined in KRT/Oligonucleotides. Droplets for droplet PCR amplification were generated using the QX200 Droplet Generator (BioRad). Data acquisition and primary analysis was done using the QX200 Droplet Reader (BioRad) and QuantaSoft software (BioRad). All steps were performed according to the manufacturer’s instructions. To calculate the number of target copies per ng RNA input, samples were normalized using *RPL30* and *RAF1* as normalization genes (Coulter, 2018). Statistical comparisons were performed in R using t-test in ggpubr package v.0.4.0 (Kassambara, 2020). Results were visualized in R using the tidyverse package (Wickham et al., 2019).

#### Global RNA-seq

The sequencing library was prepared with TruSeq Stranded mRNA Library Prep (Illumina) according to manufacturer’s instructions. The 19 libraries were pooled at equimolar concentrations and sequenced on an Illumina NovaSeq 6000 S1 flow cell (Illumina) with 100 bp paired end reads. The quality of sequencing reads was assessed using BBMap (Bushnell, 2014), and adapter sequences and low-quality reads were removed. The sequencing reads were then mapped to the GRCh38.p5 index using HISAT2 (Kim et al., 2015). Mapped paired end reads were counted to protein coding genes using featureCounts (Liao et al., 2014). Differential expression analysis was conducted in R version 3.5.1 (R Core Team, 2019) using SARTools v.1.6.8 (Varet et al., 2016) and the DESeq2 v.1.22.1 (Love et al., 2014), and genes were considered significantly differentially expressed with an FDR < 0.01. Normalized counts were visualized using the tidyverse package v.1.3.0 (Wickham et al., 2019). The heatmaps were generated using the pheatmap package version 1.0.12 (Kolde, 2019). The Wald-test was used to calculate p-values and Benjamini-Hochberg was used to correct for multiple testing. The gene ontology (GO) analysis of a ranked list of differential expressed genes were performed using GSEA software (Subramanian et al., 2005) looking at biological process (BP) terms.

#### Illumina EPIC array

DNA methylation status of 22 samples were assessed using the Infinium MethylationEPIC BeadChip v.1.0_B3 (Illumina). Quality control and pre-processing of the raw data was performed in R using Minfi v.1.36.0 (Aryee et al., 2014). No samples were removed due to poor quality (detection p values >0.05). Background correction was performed using NOOB method (Triche et al., 2013) and β values (ratio of methylated signal divided by the sum of the methylated and unmethylated signal) were normalized using functional normalization (Fortin et al., 2014). Probes with unreliable measurements (detection p values >0.01) (n = 8,818) and cross-reactive probes (Chen et al., 2013) (n = 43,256) were then removed, resulting in a final data set consisting of 814,112 probes and 22 samples. Probes were annotated with Illumina Human Methylation EPIC annotation 1.0 B5 (hg38). Differential methylation (DM) analysis was performed on the M values (log2 of the β values) using the limma package (Ritchie et al., 2015), and CpGs were considered significantly differentially methylated with an FDR < 0.01. GO analysis was performed using top 10 % DM CpGs (DMCs) as input to GOMETH in the missMethyl package version 1.24.0 (Phipson et al., 2016) for BP terms.

#### Integration of RNA-seq and DNA methylation data

Data from matching DNA and RNA samples (extracted from the same wells, n = 16) were subsetted to undergo statistical integration. Multi-Omics Regulation (MORE) (Conesa, 2018) was used to identify CpGs that regulate gene expression by applying Generalized Linear Models: normalized counts for differentially expressed genes (from DEseq2) were used as the response variable, CpG M-values (from Minfi) and experimental covariates (Day) were used as predictors. First, CpGs with low variability were filtered and multicollinearity was reduced by grouping highly correlated CpGs. Variable selection was then performed with Elastic Net regression and stepwise (two-ways backward) regression. CpGs were considered to significantly regulate gene expression when the regression coefficient p-value was < 0.05. Significant CpG regulators of gene expression were visualized using the Tidyverse package (Wickham et al., 2019) using beta values (n = 22) and normalized counts (n = 19) from all samples.

#### Collection of cells and scRNA-seq

Cells harvested on Days 0, 7, 13 and 20 were washed twice in wells with 1xPBS and detached using Accutase (STEMCELL Technologies) at 37 °C for 7 min. Cells were triturated 10-15 times to separate into single cells and transferred to centrifuge tubes containing the appropriate base media with 0.05 % BSA (Sigma-Aldrich). Counts were performed using Countess II FL Cell Counter (ThermoFisher Scientific), cells were centrifuged at 300x g for 5 min and the supernatant was discarded. Cell pellets were then resuspended in base medium containing 0.05 % BSA and cell aggregates were filtered out using MACS SmartStrainers (Miltenyi). The cells were recounted and processed within 1 hour on the 10x Chromium controller (10x Genomics). Approximately 2,300 cells were loaded per channel on the Chromium Chip B (10x Genomics) to give an estimated recovery of 1,400 cells. The Chromium Single Cell 3’ Library & Gel Bead Kit v3 (10x Genomics) and Chromium i7 Multiplex Kit (10x Genomics) were used to generate scRNA-seq libraries, according to the manufacturer’s instructions. Libraries from 16 samples were pooled together based on molarity and sequenced on a NextSeq 550 (Illumina) with 28 cycles for read 1, 8 cycles for the I7 index and 91 cycles for read 2. For the second sequencing run, libraries were pooled again based on the number of recovered cells to give a similar number of reads per cell for each sample (33,000 - 44,000 reads/cell).

#### scRNA-seq data analysis

The Cell Ranger 3.1.0 Gene Expression pipeline (10x Genomics) was used to demultiplex the raw base-call files and convert them into FASTQ files. The FASTQ files were aligned to the GRCh38-3.0.0 human reference genome, and Cell Ranger count was used with default parameters for computing read counts for Days 0, 7, 13 and 20. The sequenced replicates for each day were aggregated into single datasets using Cell Ranger Aggr command. Duplicates, dead cells and cells with greater than 5 median absolute deviations (MADs) for mitochondrial reads were filtered out (McCarthy et al., 2017). We used scTRANSFORM for normalization to better understand cell to cell heterogeneity after performing cell cycle regression analysis (Hafemeister and Satija, 2019; Tirosh et al., 2016) (for more details, see supplemental information). We used a resolution of 0.55 to cluster cells, obtained by determining the optimum number of clusters (cell grouped together sharing similar expression profiles) in the dataset using the Clustree R package (Zappia and Oshlack, 2018) (**Fig. S2B and C**).

#### scATAC-seq Library Preparation and Sequencing

Cells were washed twice with 1xPBS and detached to single cell suspension by application of Accutase (STEMCELL Technologies) at 37 °C for 7 min. The detached cells were washed with appropriate base media with added 0.04% BSA (Sigma-Aldrich) and filtered using MACS SmartStrainers (Miltenyi Biotech) to remove cell aggregates. Nuclei isolation was done according to the 10x Genomics protocol CG000169 (Rev D) using 2 minutes of incubation in lysis buffer diluted to 0.1x and 0.5x for Day 0 and Day 20 cells, respectively. We used the Countess II FL Cell Counter (ThermoFisher Scientific) to quantify nuclei and confirm complete lysis and microscopy to confirm high nuclei quality. Nuclei were further processed on the 10x Chromium controller (10x Genomics) using Next GEM Chip H Single Cell Kit (10x Genomics), Next GEM Single Cell ATAC Library & Gel Bead Kit v1.1 (10 x Genomics) and Chromium i7 Multiplex Kit N Set A (10x Genomics) according to the Next GEM Single Cell ATAC Reagent Kits v1.1 User Guide (CG000209, Rev C). The targeted nuclei recovery was 5,000 nuclei per sample. The resulting 4 sample libraries were sequenced on a NovaSeq Sp flow cell (Illumina) with 50 cycles for read 1, 8 cycles for the i7 index read, 16 cycles for the i5 index read and 49 cycles for read 2.

#### scATAC sequencing analysis

Cell Ranger ATAC version 1.2.0 with reference genome GRCh38-1.2.0 was used to pre-process scATAC-seq raw sequencing data into FASTQ files. Single cell accessibility counts for the cells were generated from reads using the ‘cellranger-atac count’ pipeline. Reference genome HG38 used for alignment and generation of single-cell accessibility counts was obtained from the 10x Genomics (https://support.10xgenomics.com/single-cell-atac/software/downloads/).

Downstream analysis of the scATAC-seq data was performed using the R package ArchR v1.0.1 (Granja et al., 2021). A tile matrix of 500-bp bins was constructed after quality control, removal of low-quality cells and doublet removal using the *doubletfinder* function of ArchR. The ArchR Project contained the filtered cells that had a TSS enrichment below 3 and <1000 fragments. A layered dimensionality reduction approach utilizing Latent Semantic Indexing (LSI) and Singular Value Decomposition (SVD) applied on Genome-wide tile matrix. Uniform Manifold approximation and projection (UMAP) was performed to visualize data in 2D space. Louvain Clustering methods implemented in R package Seurat (Stuart et al., 2019) was used for clustering of the single-cell accessibility profiles.

