## Supplemental Information for "A multi-omics approach to visualize early neuronal differentiation in 4D"

[$] Equal co-authorship contribution.

### Co-corresponding authors

[1]Division of Clinical Paediatrics, Department of Women's and Children's Health, Karolinska Institutet, Sweden
[2]Astrid Lindgren Children's Hospital Karolinska University Hospital, Stockholm, Sweden
[3]PharmaTox Strategic Research Initiative, Faculty of Mathematics and Natural Sciences, University of Oslo, Norway
[4]Department of Medical Genetics, Oslo University Hospital and University of Oslo, Norway
[5]Institute of Clinical Medicine, Faculty of Medicine, University of Oslo, Oslo, Norway.
[6]Department of Informatics, University of Oslo, Norway
[7]Institute of Basic Medical Sciences, University of Oslo, Oslo, Norway
[8]Department of Biosciences, University of Oslo, Norway
[9] Division of Obstetrics and Gynecology, Department of Clinical Science, Intervention and Technology (CLINTEC), Karolinska Institutet, Alfred Nobels Allé 8, SE-14152, Stockholm, Sweden.
[10]Center for Fetal Medicine, Karolinska University Hospital Huddinge, SE-14186 Stockholm, Sweden.
[11]Pharmacoepidemiology and Drug Safety Research Group, Department of Pharmacy, School of Pharmacy, University of Oslo, Norway
[12]Division of Clinical Neuroscience, Department of Research and Innovation, Oslo University Hospital, Oslo, Norway
[13]Centre for Fertility and Health, Norwegian Institute of Public Health, Oslo, Norway
[14]Lead contact

[*] Current address: Department of Cancer Immunology, Institute for Cancer Research, Oslo University Hospital, and KG Jebsen Centre for B-cell malignancies, Institute for Clinical Medicine, University of Oslo, Norway.

[&] Current address: Department of Medical Genetics, Oslo University Hospital, and University of Oslo, Norway.

[§] Current address: Department of Analysis and Diagnostics, Section for Molecular Biology, Norwegian Veterinary Institute, Ås, Norway.

[+] Current address: Istituto di Genetica Molecolare, CNR - Consiglio Nazionale delle Ricerche, Pavia, Italy.

#### Supplemental Figures

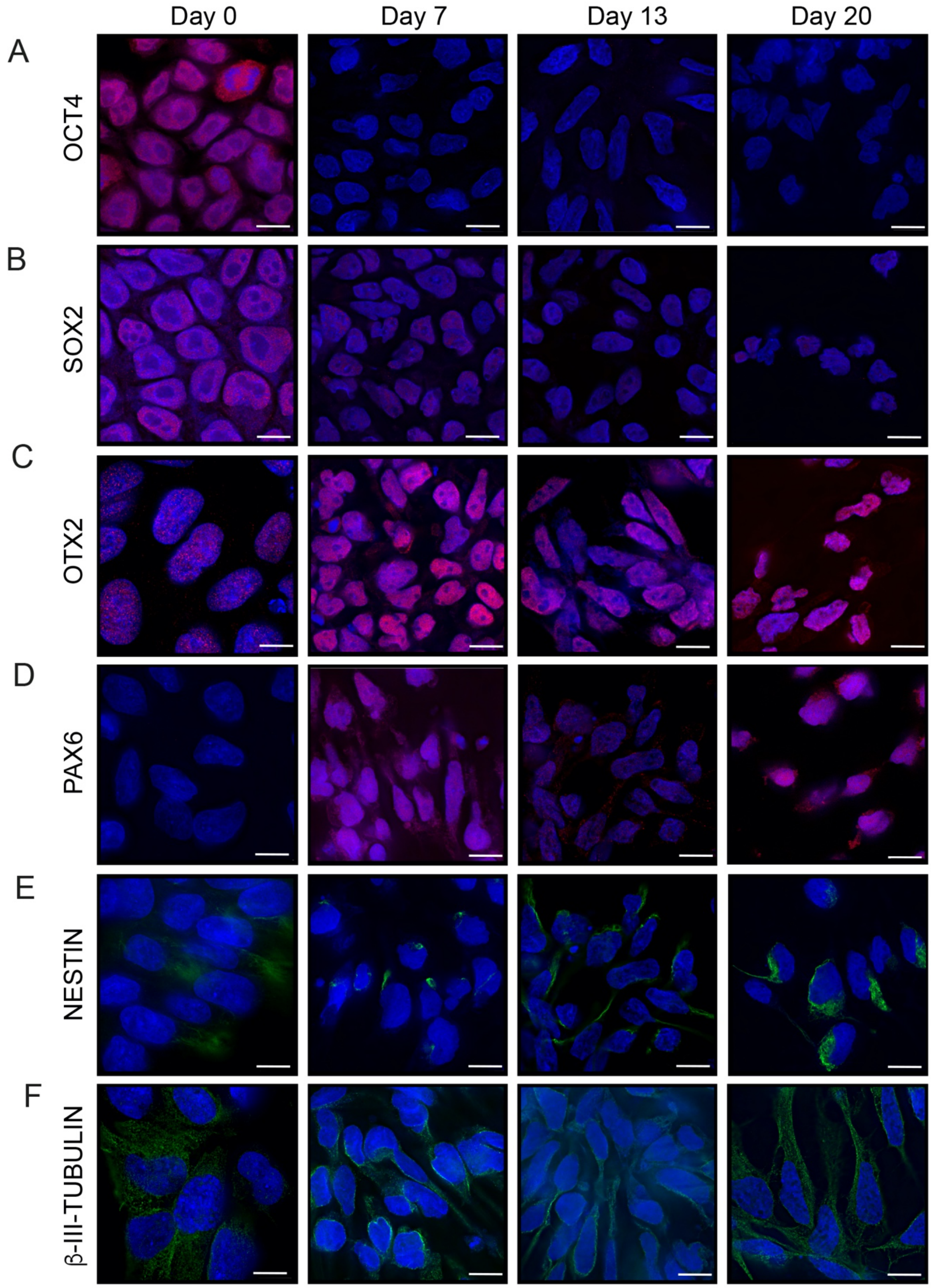


**Figure S1.** Immunofluorescence staining and imaging of pluripotency, neural and neuronal markers at Day 0, 7, 13 and 20 of differentiation, related to Figure 1. Confocal immunofluorescent micrographs show the localization of the transcription factors A) OCT4, B) SOX2, C) OTX2 and D) PAX6, in red, and of the filamentous proteins E) NESTIN and F) β-III-TUBULIN, in green. The cells were counterstained with the DNA marker DAPI (blue), and the scale bar corresponds to 5 μm.

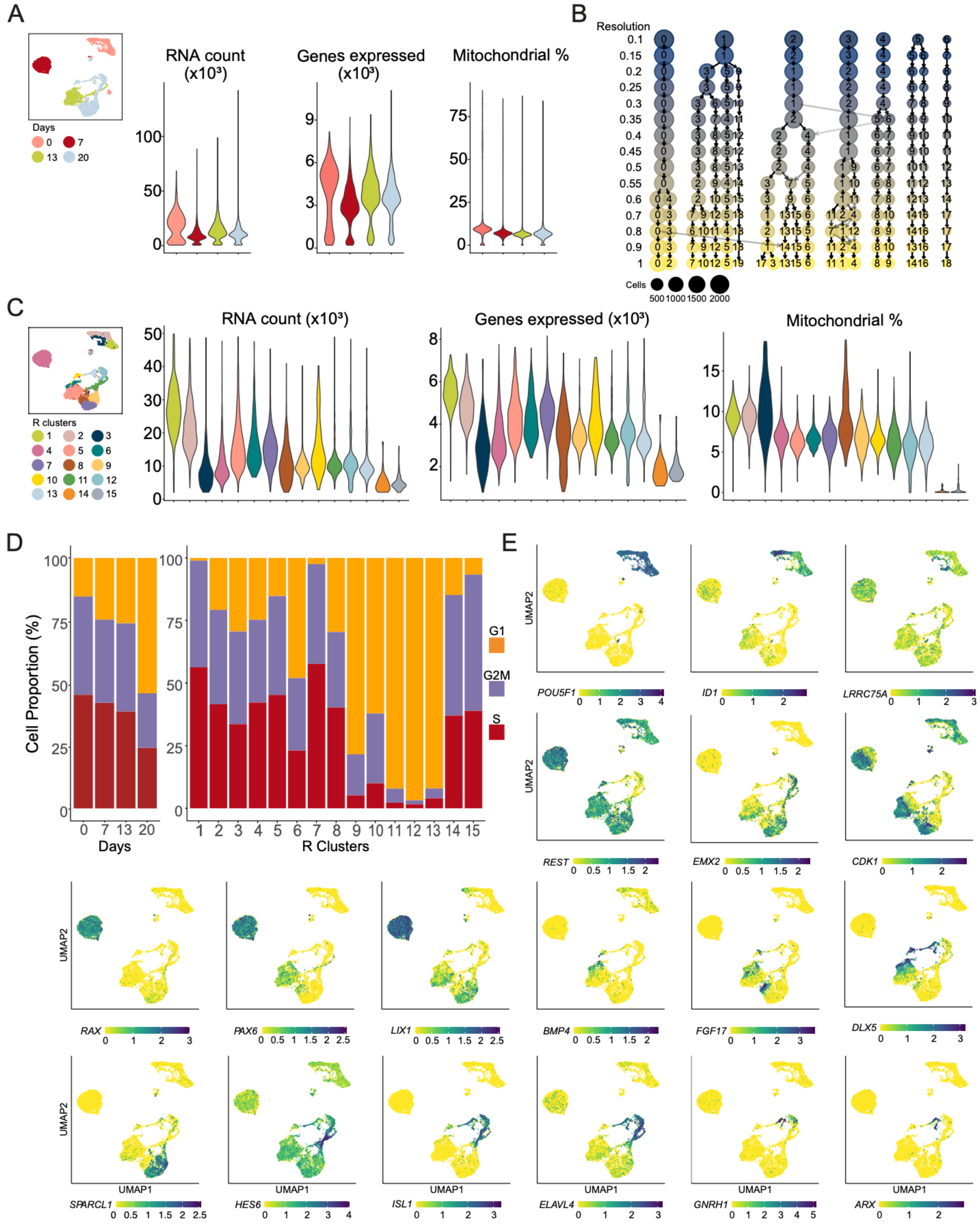


**Figure S2.** Quality control, clustering, and visualization of combined scRNA-seq datasets and genes used to profile the cell populations and annotate the clusters, related to Figure 2. A) UMAP showing original cell identity at day 0, 7, 13 and 20, and corresponding violin plots representing scRNA-seq count, number of genes expressed and mitochondrial content per cell before filtering. B) Dendrogram plot showing numbers of resulting clusters at resolutions 0.1 to 1. Circle size represents the number of cells per cluster. C) UMAP showing clusters R1-15 at resolution 0.55 and corresponding violin plots representing RNA count, number of genes expressed and mitochondrial content per cell after filtering. D) Proportion of cells in different cell cycle phases per day and per cluster at resolution 0.55. E) Representative UMAPs showing cluster specific or differentiation-driven gene expression across all four timepoints for *POU5F1, ID1, LRRC75A, REST, EMX2, CDK1, RAX, PAX6, LIX1, BMP4, FGF17, DLX5, SPARCL1, HES6, ISL1, ELAVL4, GNRH1* and *ARX.*

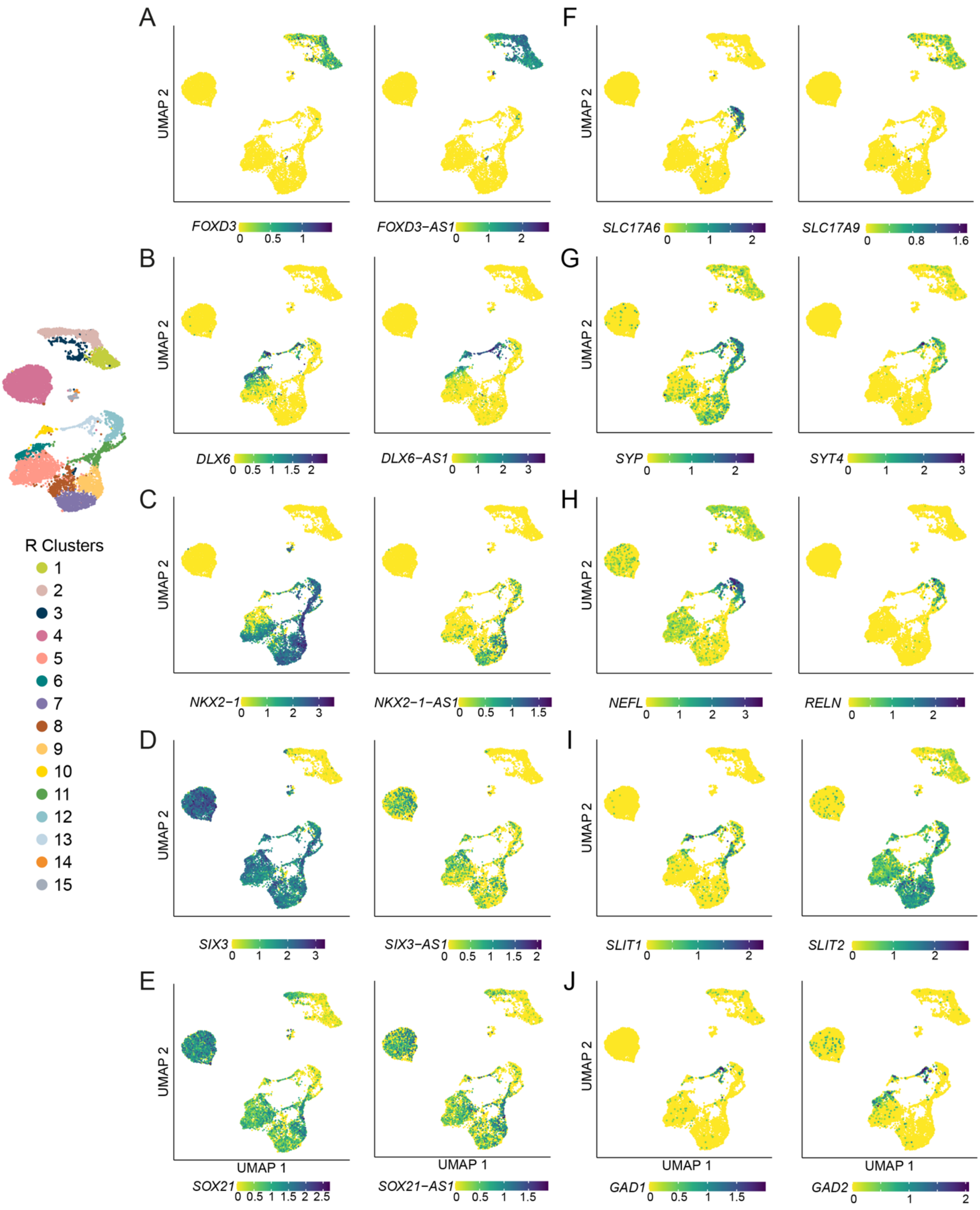


**Figure S3.** Different expression levels of antisense non-coding RNA and corresponding genes during neuronal differentiation, related to Figure 2 and 3. scRNA-seq UMAPs of individual genes and corresponding non-coding RNAs (AS1) for A) *FOXD3*, B) *DLX6*, C) *NKX2-1*, D) *SIX3* and E)

*SOX21*. F-J show selected differentiation, maturation and function related marker genes F) *SLC17A6* and *SLC17A9*, G) *SYP* and *SYT4*, H) *NEFL* and *RELN*, I) *SLIT1* and *SLIT2*, J) *GAD1* and *GAD2*. UMAP cluster annotation shown to the left.

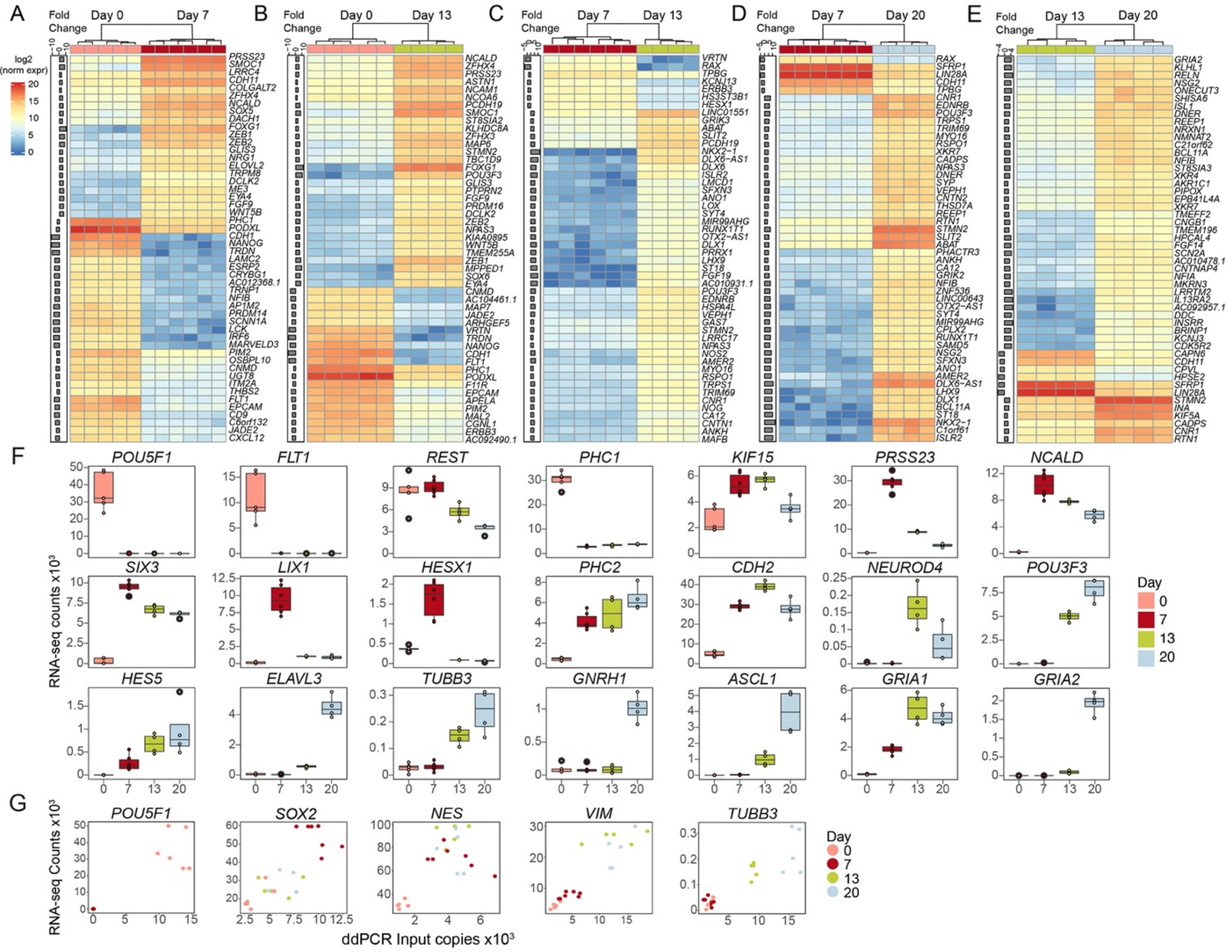


**Figure S4.** RNA-seq reveals novel marker genes during neuronal differentiation, related to Figure 4. A-E) Clustered heatmaps of the top 50 differentially expressed genes between timepoints A) Day 0 vs Day 7, B) Day 0 vs Day 13, C) Day 7 vs Day 13, D) Day 7 vs Day 20 and E) Day 13 vs Day 20. Fold change is shown to the left. F) Expression pattern of genes *POU5F1, FLT1, REST, PHC1, KIF15, PRSS23, NCALD, SIX3, LIX1, HESX1, PHC2, CDH2, NEUROD4, POU3F3, HES5, ELAVL3, TUBB3, GNRH1, ASCL1, GRIA1,* and *GRIA2* across differentiation. G) Correlation between read counts of the RNA-seq data and ddPCR input copies as shown in the scatter plots for marker genes *POU5F1, SOX2, NES, VIM* and *TUBB3*. Datapoints are color-coded in pink for Day 0, red for Day 7, green for Day 13 and blue for Day 20 in both G and H.

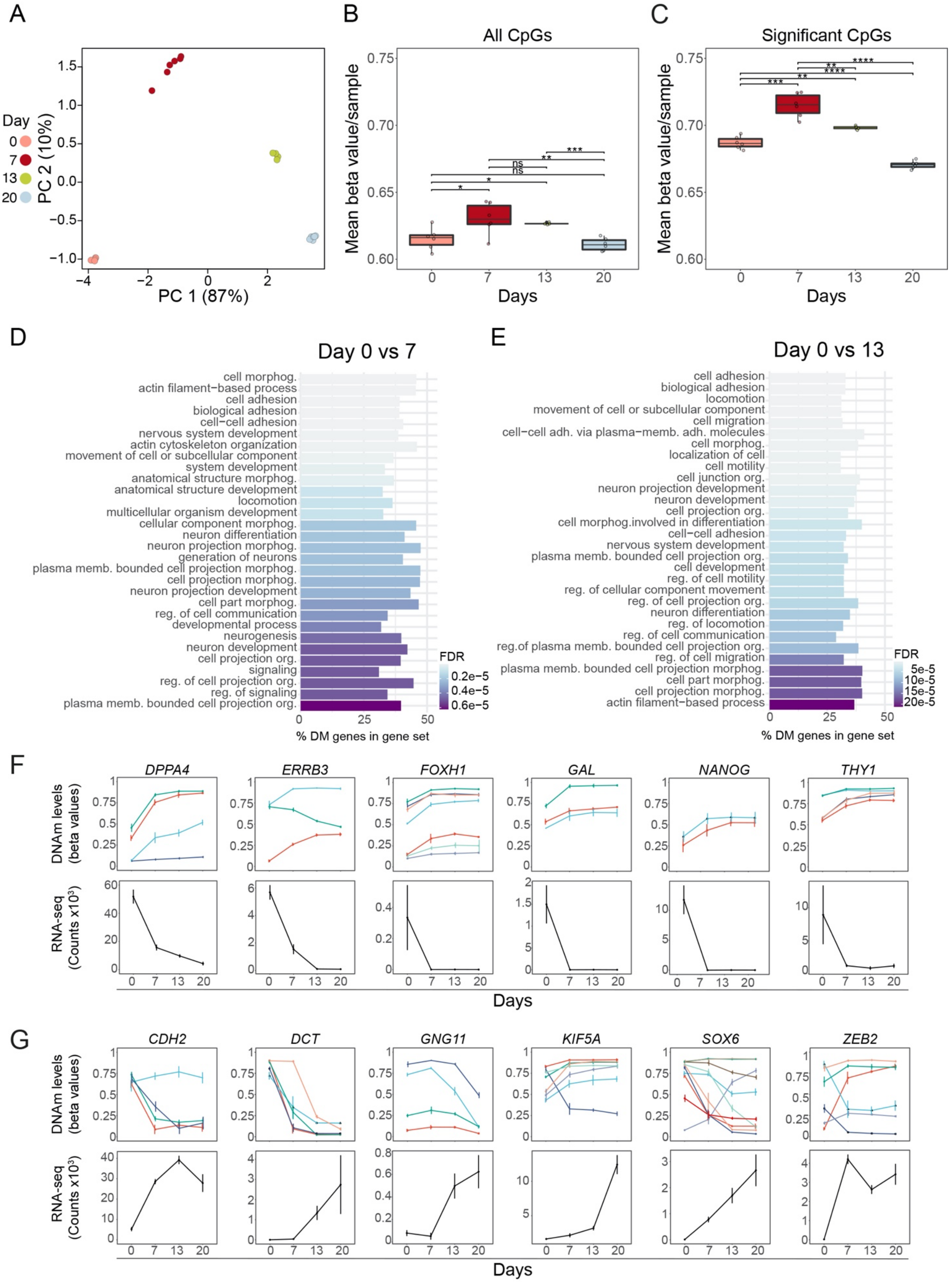


**Figure S5.** Genome-wide DNA methylation status changes during neuronal differentiation, related to Figure 5. A) Principal component analysis for all replicates for Days 0, 7, 13 and 20. B) Mean DNAm levels (beta-values) per sample for all CpGs across differentiation for Days 0, 7, 13 and 20. C) Mean DNAm levels (beta-values) per sample for all significant CpGs at Days 0, 7, 13 and 20. B-

C) *p < 0.05, **p < 0.01, ***p < 0.001, ****p < 0.0001. D-E) Top 30 ranked over-represented gene ontology (GO) terms in biological process (BP) (GO-BP) on the top 10% differentially methylated CpGs, regarding transition from Day 0 to Day 7 (D) and Day 0 to Day 13 (E). F) Examples of inverse correlation between DNAm and gene expression levels across neuronal differentiation, at the level of significant CpG regulators, as derived by the MORE analysis. For genes becoming repressed during differentiation, such as *DPPA4, ERRB3, FOXH1, GAL, NANOG* and *THY1,* DNAm levels increased as seen for some significant CpGs. Top panels show mean +/- standard deviation of DNAm and bottom panels show normalized RNA-seq counts. (G) MORE analysis of *CDH2, DCT, GNG11, KIF5A, SOX6* and *ZEB2,* with DNAm levels decreasing upon transcriptional activation during differentiation. Top panels show mean +/- standard deviation of DNAm and bottom panels show normalized RNA-seq counts.

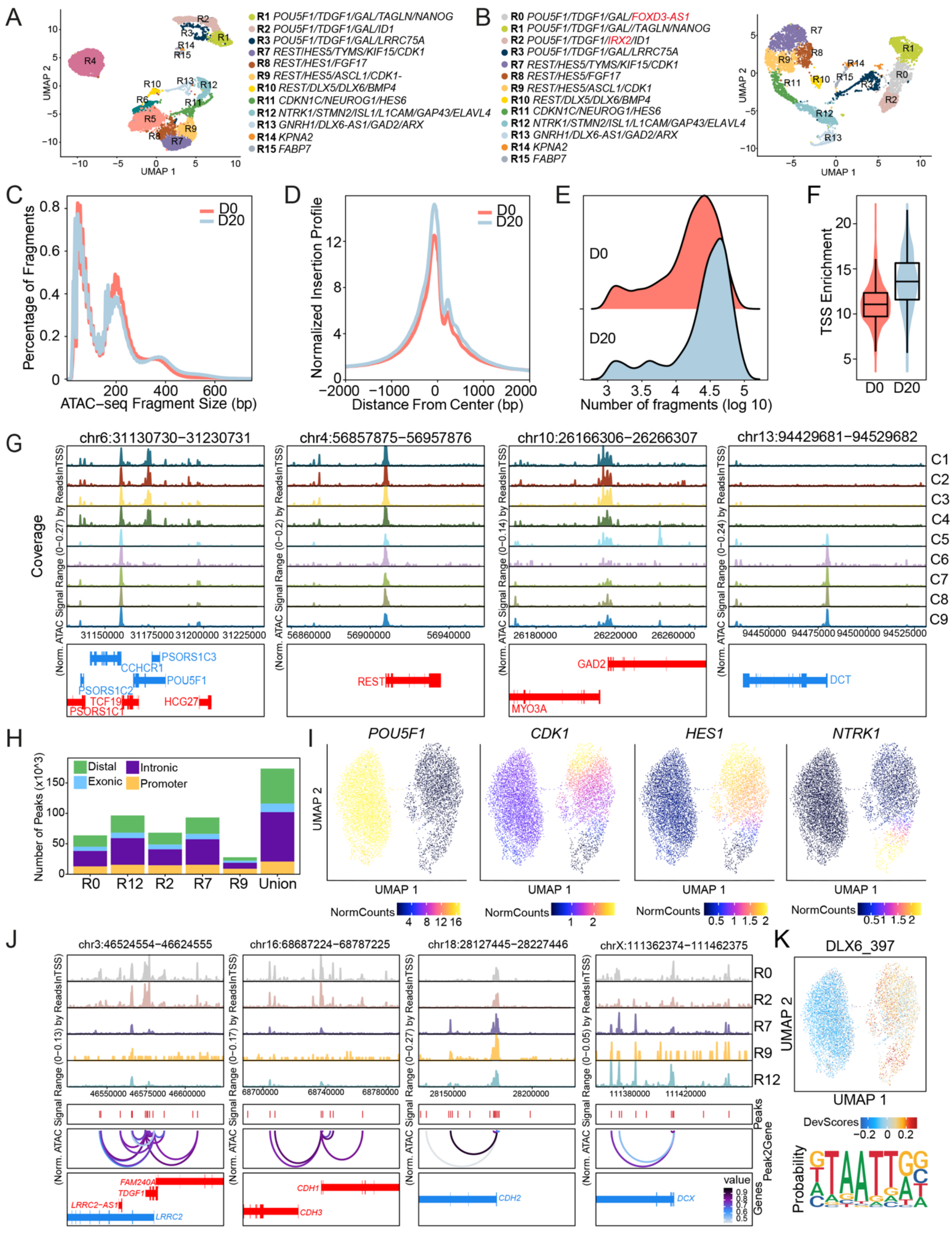


**Figure S6.** Gene expression and chromatin opening features during neuronal differentiation, related to Figure 6. A) Single-cell UMAP plot and cluster annotations for all timepoints Day 0, 7, 13 and 20. B) Single-cell UMAP plot and cluster annotations for timepoints Day 0 and 20. New cluster marker genes in red font. C) Percentage of ATAC-seq fragments and corresponding fragment size in base pairs for Day 0 and 20. D) Normalized insertion profiles and corresponding distance from centre in base pairs for Day 0 and 20. E) Number of fragments and F) TSS distribution of scATAC-seq datasets represented as a box plot; where the middle represents the median, and the lower value is 25th percentile and the upper hinge is the 75th percentile of the data. Interquartile range (IQR) represent the distance between the upper and lower hinges and the whisker represents represent the lowest and largest values within 1.5 times the IQR. G) Genome track visualization of ATAC-seq per cluster for *POU5F1, REST, GAD2, CDH2* and *DCT* gene loci. H) Distribution of ATAC-seq peaks per cluster at promoter, intronic, exonic and distal regions for integrated clusters R0 (n=5027), R2 (n=448), R7 (n=2343), R9 (n=145), R12 (n=552), and all union peaks. I) UMAPs representing Gene Integration Matrix for *POU5F1, CDK1, HES1* and *NTRK1.* J) Genome tracks of ATAC-seq peaks in integrated clusters R0, R2, R7, R9 and R12 for *TDGF1, CDH1, CDH2* and *DCX* loci. Peaks and inferred peak-to-gene links for distal regulatory elements across the differentiation dataset are shown below. K) Motif matrix UMAP for DLX6 with corresponding representative sequence logo.

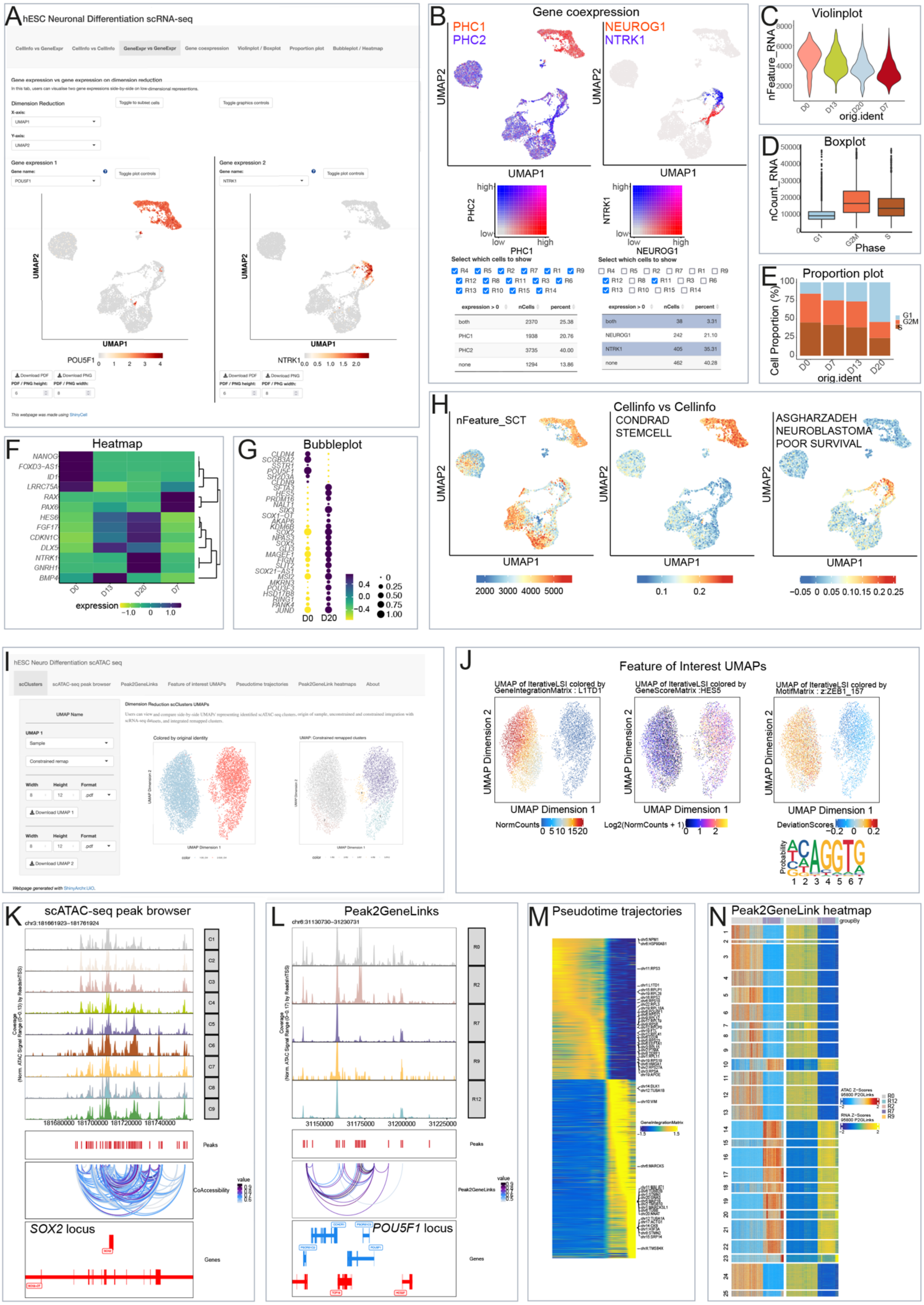

**Figure S7.** Visualizing gene expression and chromatin openness during neuronal differentiation at single-cell resolution in open access web interfaces, related to Figure 6, S6 and 7.

A) Visualization of the scRNA-seq data utilizing the ShinyCell tool. The user can explore the data in a web interface with seven different tabs. Gene expression can be explored for genes of interest in a tab together with cell information or in a gene expression tab as two side-by-side UMAPs, here shown for *POU5F1* and *NTRK1*. B) The gene co-expression tab allows inspection of overlapping expression of two genes in all clusters or selected clusters with downloadable number of cells and percentage data. Here illustrated for two gene pairs: *PHC1/PHC2*, and *NEUROG1/NTRK1* C) Violin plots, D) box plots and E) proportion plots can be generated to explore a number of features including cell cycle analysis and RNA count. The expression of a list of selected genes can be further explored by F) Heatmap or G) Bubble plot. H) In the cell info tab, the user can explore cellular features such as clusters, heterogeneity in the data by for example single-cell transcriptomics (SCT) and pathway analysis. We present another web interface that allows for exploration of chromatin openness utilizing our in-lab ShinyArchR.UiO tool. I) The user can analyse the scATAC-seq data in six different tabs. In the scClusters tab, the user can visualize scATAC-seq data including original sample and cluster UMAPs. J) The feature of interest tab allows for exploration of gene score matrix, gene integration matrix and motif matrix UMAPs of selected genes, here shown for *L1TD1, HES5* and *ZEB1*. K) The user can also explore scATAC-seq clusters and integrated clusters in genome-browser views for selected gene loci with peaks and co-accessibility, here with an example from the *SOX2* locus. L) ATAC-seq peaks can also be explored across integrated clusters, as shown for the *POU5F1* locus. Finally, two tabs display different heatmaps including M) gene integration matrix as shown, but also gene score matrix, motif matrix and peak matrix can be explored side-by side. N) In the peak2genelink tab, a heatmap of gene scores from scATAC-seq (to the left) with complementary gene expression from scRNA-seq data (to the right) are grouped by integrated clusters represented by coloured of bars on top. All plots can be downloaded as high-resolution PNGs or PDFs.

### Supplemental tables

**Table S1.** Cell Ranger QC output related to Figure 2.

| Day | Replicate | Estimated cell number | Median UMIs/cell | Median genes/cell | Mean reads/cell | Sequencing saturation (%) |
|---|---|---|---|---|---|---|
| 0 | 1 | 1129 | 19501 | 4719 | 39466 | 11.1 |
| 0 | 2 | 1360 | 22989 | 5082 | 44124 | 13.4 |
| 7 | 1 | 1477 | 13610 | 4142 | 38397 | 18.3 |
| 7 | 2 | 1385 | 13231 | 3417 | 38248 | 20.9 |
| 13 | 1 | 977 | 13931 | 4100 | 41081 | 19.5 |
| 13 | 2 | 1409 | 13212 | 3864 | 36816 | 19.1 |
| 20 | 1 | 1816 | 9892 | 3485 | 34531 | 29.1 |
| 20 | 2 | 1614 | 11055 | 3711 | 36155 | 28.1 |

**Table S2.** Cell counts and populations (%) per cell-cycle phase at the stem cell level and the three distinct timepoints of the neuronal differentiation protocol, i.e., Day 7, 13 and 20, related to Figure 2 and S2.

| Timepoint | Cell Cycle Phase Cell counts (%) | | |
|---|---|---|---|
| | G1 | G2M | S |
| Day 0 | 292 (15.4) | 747 (39.3) | 861 (45.3) |
| Day 7 | 584 (24.7) | 785 (33.2) | 999 (42.2) |
| Day 13 | 533 (26.1) | 721 (35.3) | 791 (38.7) |
| Day 20 | 1635 (54.1) | 657 (21.7) | 732 (24.2) |

**Table S3.** Cell numbers and cluster populations (%) per timepoint, related to Figure 2.

| | Day 0 | Day 7 | Day 13 | Day 20 | Total |
|---|---|---|---|---|---|
| **R1** | 636 (33.5) | 0 (0) | 0 (0) | 0 (0) | 636 |
| **R2** | 953 (50.2) | 0 (0) | 0 (0) | 0 (0) | 953 |
| **R3** | 283 (14.9) | 1 (0.04) | 1 (0.1) | 4 (0.1) | 289 |
| **R4** | 0 (0) | 2341 (98.9) | 3 (0.1) | 0 (0) | 2344 |
| **R5** | 0 (0) | 0 (0) | 1512 (73.9) | 47 (1.6) | 1559 |
| **R6** | 0 (0) | 0 (0) | 273 (13.3) | 0 (0) | 273 |
| **R7** | 0 (0) | 0 (0) | 2 (0.1) | 851 (28.1) | 853 |
| **R8** | 1 (0.1) | 7 (0.3) | 36 (1.8) | 361 (11.9) | 405 |
| **R9** | 0 (0) | 0 (0) | 2 (0.1) | 619 (20.5) | 621 |
| **R10** | 0 (0) | 0 (0) | 0 (0) | 140 (4.6) | 140 |
| **R11** | 0 (0) | 1 (0.04) | 164 (8.0) | 230 (7.6) | 395 |
| **R12** | 0 (0) | 0 (0) | 4 (0.2) | 495 (16.4) | 499 |
| **R13** | 0 (0) | 0 (0) | 29 (1.4) | 224 (7.4) | 253 |
| **R14** | 27 (1.4) | 0 (0) | 0 (0) | 0 (0) | 27 |
| **R15** | 0 (0) | 18 (0.8) | 19 (0.9) | 53 (1.8) | 90 |

**Table S4.** Global gene expression and DNA methylation changes between timepoints, related to Figures 4, S4, 5, and S5.*

| | Day 0 vs day 7 | Day 0 vs day 13 | Day 0 vs day 20 | Day 7 vs day 13 | Day 7 vs day 20 | Day 13 vs day 20 |
|---|---|---|---|---|---|---|
| # DE genes (down, up) | 8972 (5025, 3946) | 9252 (4948, 4304) | 11313 (5710, 5603) | 3602 (1624, 1978) | 7120 (3258, 3861) | 2379 (1529, 2084) |
| # DM CpGs (down, up) | 161600 (24528, 137132) | 146870 (46130, 100740) | 210049 (94265, 115784) | 39545 (31372, 8173) | 122781 (105161, 17620) | 47676 (45727, 1949) |
| # DM CpGs annotated to genes | 110994 | 100816 | 143763 | 25337 | 79647 | 31082 |
| # DMG overlapping with DEG (% of DEGs) | 6446 (72) | 6639 (72) | 8388 (74) | 1862 (52) | 5117 (72) | 1414 (59) |
| # DE genes regulated by CpGs (MORE) | 6441 | 6775 | 8011 | 2683 | 5230 | 1870 |
| # Significant CpG regulators (MORE) | 16815 | 18950 | 21199 | 8072 | 14542 | 5651 |

*The number of differentially expressed genes (DEG) and differentially methylated (DM) CpGs are described for all comparisons. Of the DM CpGs that are annotated to genes, the overlap with DEG varies from 48 - 73%. The Multi-Omics regulations (MORE) analysis reveals the number of DEG that are regulated by CpGs and the number of CpG regulators.

**Table S5.** GSEA for all timepoints in the protocol, related to Figure 4 and S4.

| Comparison | GO term | NES | FDR |
|---|---|---|---|
| Day 0 vs day 7 | DORSAL VENTRAL PATTERN FORMATION | 1,892 | 0,024 |
| | PALLIUM DEVELOPMENT | 1,886 | 0,015 |
| | EAR MORPHOGENESIS | 1,884 | 0,010 |
| | PATTERN SPECIFICATION PROCESS | 1,881 | 0,009 |
| | INNER EAR MORPHOGENESIS | 1,869 | 0,011 |
| | NEURON FATE COMMITMENT | 1,858 | 0,013 |
| | NEUROBLAST PROLIFERATION | 1,845 | 0,016 |
| | MICROTUBULE BUNDLE FORMATION | 1,844 | 0,015 |
| | SENSORY ORGAN MORPHOGENESIS | 1,835 | 0,017 |
| | LENS DEVELOPMENT IN CAMERA TYPE EYE | 1,834 | 0,016 |
| | CEREBRAL CORTEX DEVELOPMENT | 1,824 | 0,019 |
| | POSITIVE REGULATION OF NEURAL PRECURSOR CELL PROLIFERATION | 1,818 | 0,020 |
| | CELL FATE DETERMINATION | 1,801 | 0,030 |
| | REGULATION OF NEUROBLAST PROLIFERATION | 1,790 | 0,036 |
| | DEVELOPMENTAL INDUCTION | 1,788 | 0,035 |
| | CELL AGGREGATION | 1,773 | 0,047 |
| | AXONEME ASSEMBLY | 1,769 | 0,049 |
| | POSITIVE REGULATION OF CARDIOCYTE DIFFERENTIATION | 1,768 | 0,046 |
| | POSITIVE REGULATION OF NEUROBLAST PROLIFERATION | 1,767 | 0,045 |
| | FOREBRAIN REGIONALIZATION | 1,766 | 0,044 |
| Day 0 vs day 13 | PALLIUM DEVELOPMENT | 2,024 | < 0,00001 |
| | TELENCEPHALON DEVELOPMENT | 1,955 | 0,0003 |
| | DORSAL VENTRAL PATTERN FORMATION | 1,949 | 0,0003 |
| | PATTERN SPECIFICATION PROCESS | 1,945 | 0,0003 |
| | FOREBRAIN GENERATION OF NEURONS | 1,929 | 0,0004 |
| | NEURON MIGRATION | 1,928 | 0,0004 |
| | MICROTUBULE BUNDLE FORMATION | 1,918 | 0,0005 |
| | AXONEME ASSEMBLY | 1,915 | 0,0006 |
| | FOREBRAIN CELL MIGRATION | 1,904 | 0,0008 |
| | CEREBRAL CORTEX DEVELOPMENT | 1,902 | 0,0007 |
| | SENSORY ORGAN MORPHOGENESIS | 1,899 | 0,0008 |
| | CILIUM ORGANIZATION | 1,898 | 0,0007 |
| | EAR MORPHOGENESIS | 1,898 | 0,0007 |
| | HIPPOCAMPUS DEVELOPMENT | 1,895 | 0,0007 |
| | NEURON FATE COMMITMENT | 1,884 | 0,0009 |
| | POSITIVE REGULATION OF NEURAL PRECURSOR CELL PROLIFERATION | 1,881 | 0,0010 |
| | CENTRAL NERVOUS SYSTEM NEURON DEVELOPMENT | 1,880 | 0,0009 |
| | CEREBRAL CORTEX CELL MIGRATION | 1,875 | 0,001 |
| | CENTRAL NERVOUS SYSTEM NEURON DIFFERENTIATION | 1,863 | 0,001 |
| | NEUROBLAST PROLIFERATION | 1,857 | 0,002 |
| | EMBRYONIC ORGAN MORPHOGENESIS | 1,853 | 0,002 |
| | INNER EAR MORPHOGENESIS | 1,851 | 0,002 |
| | FOREBRAIN NEURON DIFFERENTIATION | 1,849 | 0,002 |
| | KERATINIZATION | -1,844 | 0,022 |
| | FOREBRAIN DEVELOPMENT | 1,837 | 0,003 |
| | REGIONALIZATION | 1,836 | 0,003 |
| | APICAL JUNCTION ASSEMBLY | -1,832 | 0,018 |
| | CELL FATE DETERMINATION | 1,825 | 0,004 |
| | NEURAL PRECURSOR CELL PROLIFERATION | 1,825 | 0,003 |
| | EMBRYONIC SKELETAL SYSTEM DEVELOPMENT | 1,823 | 0,003 |
| | NCRNA PROCESSING | -2,060 | 0,017 |
| | REGULATION OF NEUROTRANSMITTER TRANSPORT | 1,827 | 0,001 |
| | NEUROTRANSMITTER SECRETION | 1,820 | 0,001 |
| | LOCOMOTORY BEHAVIOR | 1,807 | 0,001 |
| | NEUROTRANSMITTER TRANSPORT | 1,794 | 0,002 |
| | REGULATION OF NEUROTRANSMITTER LEVELS | 1,792 | 0,001 |
| | TRANSMISSION OF NERVE IMPULSE | 1,779 | 0,002 |
| | REGULATION OF SYNAPTIC VESICLE CYCLE | 1,775 | 0,003 |
| | NEURON MIGRATION | 1,768 | 0,003 |
| | SYNAPTIC VESICLE EXOCYTOSIS | 1,763 | 0,003 |
| | REGULATION OF SYNAPTIC PLASTICITY | 1,761 | 0,003 |
| | VESICLE MEDIATED TRANSPORT IN SYNAPSE | 1,760 | 0,003 |
| | REGULATION OF SYNAPTIC VESICLE EXOCYTOSIS | 1,753 | 0,004 |
| | NEURAL NUCLEUS DEVELOPMENT | 1,750 | 0,005 |

| | | | |
|---|---|---|---|
| Day 7 vs day 20 | SENSORY PERCEPTION OF PAIN | 1,740 | 0,007 |
| | HIPPOCAMPUS DEVELOPMENT | 1,730 | 0,010 |
| | AMINE TRANSPORT | 1,729 | 0,009 |
| | DOPAMINE TRANSPORT | 1,728 | 0,009 |
| | REGULATION OF NEURONAL SYNAPTIC PLASTICITY | 1,726 | 0,010 |
| | REGULATION OF DOPAMINE SECRETION | 1,721 | 0,011 |
| | RESPONSE TO NICOTINE | 1,711 | 0,015 |
| | EXCRETION | 1,709 | 0,016 |
| | MEMORY | 1,708 | 0,015 |
| | BEHAVIOR | 1,706 | 0,016 |
| | RESPONSE TO ETHANOL | 1,705 | 0,016 |
| | REGULATION OF ACUTE INFLAMMATORY RESPONSE | 1,700 | 0,018 |
| | NEURON MATURATION | 1,693 | 0,022 |
| | MONOAMINE TRANSPORT | 1,691 | 0,023 |
| | CATECHOLAMINE SECRETION | 1,688 | 0,024 |
| | REGULATION OF EXOCYTOSIS | 1,684 | 0,027 |
| Day 13 vs day 20 | SOMATIC STEM CELL POPULATION MAINTENANCE | -2,290 | 0,004 |
| | MAINTENANCE OF CELL NUMBER | -2,281 | 0,004 |
| | PRODUCTION OF SMALL RNA INVOLVED IN GENE SILENCING BY RNA | -2,258 | 0,005 |
| | NCRNA PROCESSING | -2,247 | 0,005 |
| | REGULATION OF POSTTRANSCRIPTIONAL GENE SILENCING | -2,233 | 0,007 |
| | SISTER CHROMATID SEGREGATION | -2,209 | 0,013 |
| | REGULATION OF GENE SILENCING | -2,207 | 0,011 |
| | MITOTIC SISTER CHROMATID SEGREGATION | -2,190 | 0,017 |
| | DIGESTIVE TRACT MORPHOGENESIS | -2,187 | 0,016 |
| | NEURAL TUBE FORMATION | -2,136 | 0,048 |
| | RIBOSOME BIOGENESIS | -2,136 | 0,044 |
| | CHROMOSOME SEGREGATION | -2,132 | 0,045 |
| | NEUROTRANSMITTER TRANSPORT | 2,118 | < 0,00001 |
| | NEUROTRANSMITTER SECRETION | 2,094 | < 0,00001 |
| | REGULATION OF NEUROTRANSMITTER LEVELS | 2,055 | < 0,00001 |
| | VESICLE MEDIATED TRANSPORT IN SYNAPSE | 2,053 | < 0,00001 |
| | REGULATION OF NEUROTRANSMITTER TRANSPORT | 2,050 | < 0,00001 |
| | REGULATION OF SYNAPTIC PLASTICITY | 2,048 | < 0,00001 |
| | SYNAPTIC VESICLE EXOCYTOSIS | 2,018 | < 0,00001 |
| | REGULATION OF SYNAPTIC VESICLE CYCLE | 1,999 | < 0,00001 |
| | LOCOMOTORY BEHAVIOR | 1,967 | < 0,00001 |
| | REGULATION OF SYNAPTIC VESICLE EXOCYTOSIS | 1,966 | < 0,00001 |
| | SYNAPTIC SIGNALING | 1,953 | 0,00001 |
| | REGULATION OF POSTSYNAPTIC MEMBRANE POTENTIAL | 1,953 | 0,00001 |
| | REGULATION OF REGULATED SECRETORY PATHWAY | 1,930 | 0,00009 |
| | REGULATION OF TRANS SYNAPTIC SIGNALING | 1,925 | 0,00014 |
| | AMINE TRANSPORT | 1,922 | 0,00014 |
| | POSITIVE REGULATION OF SYNAPTIC TRANSMISSION | 1,920 | 0,00016 |
| | SYNAPSE ORGANIZATION | 1,917 | 0,00017 |
| | RESPONSE TO NICOTINE | 1,900 | 0,00037 |

**Table S6.** Overview of significant CpG regulators of genes, related to Figures 5D and S5F-G.

| Gene | Significant CpG regulator | p-value |
|---|---|---|
| *POU5F1* | cg19865159 | 2,50E-03 |
| | cg00357273 | 1,30E-10 |
| | cg04099091 | 1,30E-10 |
| | cg09031041 | 1,30E-10 |
| | cg13083810 | 1,30E-10 |
| | cg15278102 | 1,30E-10 |
| | cg15948871 | 1,30E-10 |
| | cg16626479 | 1,30E-10 |
| | cg17364250 | 1,30E-10 |
| *NR6A1* | cg06321925 | 5,30E-06 |
| | cg08224390 | 5,30E-06 |
| | cg13539372 | 5,30E-06 |
| | cg15096615 | 5,30E-06 |
| | cg16717651 | 5,30E-06 |
| | cg12954102 | 2,05E-05 |
| | cg14156446 | 2,05E-05 |
| *DNMT3B* | cg01281600 | 1.30E-10 |
| | cg09135144 | 1.30E-10 |
| | cg09835408 | 1.30E-10 |
| | cg14224313 | 1.30E-10 |
| | cg17475857 | 1.30E-10 |
| | cg17482740 | 1.30E-10 |
| | cg17697897 | 1.30E-10 |
| | cg19184414 | 1.30E-10 |
| | cg22052056 | 1.30E-10 |
| | cg22605822 | 1.30E-10 |
| | cg24403338 | 1.30E-10 |
| | cg26553763 | 1.30E-10 |
| *CDH1* | cg07762788 | 2,60E-02 |
| | cg05303053 | 1,30E-02 |
| | cg00632260 | 1,00E-02 |
| | cg08051386 | 1,40E-03 |
| | cg13947099 | 1,40E-03 |
| | cg15653892 | 1,40E-03 |
| | cg19834745 | 1,40E-03 |
| | cg20750889 | 1,40E-03 |
| | cg23166227 | 1,40E-03 |
| | cg25218831 | 1,40E-03 |
| | cg26508465 | 1,40E-03 |
| *LDHA* | cg01316516 | 7.34E-11 |

| | | |
|---|---|---|
| | cg01381301 | 7.34E-11 |
| | cg02232751 | 7.34E-11 |
| | cg03177631 | 7.34E-11 |
| | cg03628148 | 7.34E-11 |
| | cg07187103 | 7.34E-11 |
| | cg08861826 | 7.34E-11 |
| | cg09436562 | 7.34E-11 |
| | cg11166108 | 7.34E-11 |
| | cg13190306 | 7.34E-11 |
| | cg15700009 | 7.34E-11 |
| | cg19316551 | 7.34E-11 |
| | cg19631472 | 7.34E-11 |
| | cg20429911 | 7.34E-11 |
| *KRT18* | cg11722496 | 8.56E-02 |
| | cg16747714 | 3.28E-02 |
| | cg06024289 | 2.97E-05 |
| | cg07051221 | 2.97E-05 |
| | cg07143715 | 2.97E-05 |
| | cg07652628 | 2.97E-05 |
| | cg12121782 | 2.97E-05 |
| | cg14212748 | 2.97E-05 |
| | cg27502493 | 2.97E-05 |
| *NCAM1* | cg14673319 | 3,30E-04 |
| | cg14971026 | 1,90E-03 |
| | cg00539883 | 3,30E-02 |
| | cg02804231 | 3,30E-02 |
| | cg08743050 | 3,30E-02 |
| | cg12349185 | 3,30E-02 |
| | cg13463897 | 3,30E-02 |
| | cg19624354 | 3,30E-02 |
| | cg20857767 | 3,30E-02 |
| *GAP43* | cg03395990 | 7,20E-03 |
| | cg00866813 | 2,20E-02 |
| | cg01710607 | 2,20E-02 |
| | cg04854673 | 2,20E-02 |
| | cg15143702 | 2,20E-02 |
| *CDKN1C* | cg00374747 | 2,60E-04 |
| | cg05267118 | 2,60E-04 |
| | cg10929140 | 2,80E-03 |
| | cg11399776 | 2,80E-03 |
| | cg23225147 | 2,60E-04 |
| | cg23904595 | 2,60E-04 |

| | | |
|---|---|---|
| *DCX* | cg15334207 | 8,80E-05 |
| | cg27175910 | 4,30E-02 |
| | cg11031647 | 8,90E-05 |
| | cg13936874 | 8,90E-05 |
| *ZIC4* | cg04079301 | 1.28E-03 |
| | cg20939084 | 5.56E-03 |
| | cg00896370 | 2.37E-03 |
| | cg02820514 | 2.37E-03 |
| | cg08393041 | 2.37E-03 |
| | cg13897134 | 2.37E-03 |
| | cg15105326 | 2.37E-03 |
| | cg16768018 | 2.37E-03 |
| | cg16790847 | 2.37E-03 |
| | cg17569743 | 2.37E-03 |
| | cg25449440 | 2.37E-03 |
| | cg26386385 | 2.37E-03 |
| | cg26790247 | 2.37E-03 |
| *LHX2* | cg19264651 | 1.27E-02 |
| | cg00982919 | 9.21E-03 |
| | cg03306024 | 3.59E-04 |
| | cg03559209 | 3.59E-04 |
| | cg03709033 | 3.59E-04 |
| | cg04094300 | 3.59E-04 |
| | cg05777357 | 3.59E-04 |
| | cg05900009 | 3.59E-04 |
| | cg06890720 | 3.59E-04 |
| | cg12002589 | 3.59E-04 |
| | cg13342881 | 3.59E-04 |
| | cg13585776 | 3.59E-04 |
| | cg14071579 | 3.59E-04 |
| | cg14425564 | 3.59E-04 |
| | cg16427654 | 3.59E-04 |
| | cg20939662 | 3.59E-04 |
| | cg20955788 | 3.59E-04 |
| | cg21195468 | 3.59E-04 |
| *DPPA4* | cg19250607 | 1,80E-05 |
| | cg12443195 | 4.69E-09 |
| | cg13358761 | 4.69E-09 |
| | cg18560366 | 4.69E-09 |
| *ERRB3* | cg27488983 | 1,20E-04 |
| | cg00907267 | 8,20E-03 |
| | cg26344379 | 8,20E-03 |

| | | |
|---|---|---|
| *FOXH1* | cg24947777 | 2,40E-03 |
| | cg03667317 | 1,20E-03 |
| | cg04515567 | 1,20E-03 |
| | cg07044391 | 1,20E-03 |
| | cg07205791 | 1,20E-03 |
| | cg13159060 | 1,20E-03 |
| | cg21293425 | 1,20E-03 |
| *GAL* | cg22007237 | 6,50E-03 |
| | cg01545145 | 4,50E-02 |
| | cg23009355 | 4,50E-02 |
| *NANOG* | cg10660253 | 3,90E-02 |
| | cg18781032 | 2,40E-03 |
| *THY1* | cg14651687 | 1,00E-03 |
| | cg13248115 | 2,70E-02 |
| | cg14850728 | 2,70E-02 |
| | cg15449664 | 2,70E-02 |
| | cg21633698 | 2,70E-02 |
| *CDH2* | cg18666853 | 3,40E-04 |
| | cg15322562 | 2,10E-02 |
| | cg23244545 | 2,90E-02 |
| | cg15859602 | 6,00E-04 |
| *DCT* | cg18910635 | 1.28E-07 |
| | cg05256043 | 4.37E-05 |
| | cg05807291 | 4.37E-05 |
| | cg09189889 | 4.37E-05 |
| | cg18293281 | 4.37E-05 |
| *GNG11* | cg00536924 | 3,90E-07 |
| | cg08038054 | 3,90E-07 |
| | cg08236022 | 3,90E-07 |
| | cg25546811 | 3,90E-07 |
| *KIF5A* | cg05024573 | 2,34E-05 |
| | cg06667339 | 2,34E-05 |
| | cg16949674 | 2,34E-05 |
| | cg17083982 | 2,34E-05 |
| | cg18628483 | 2,34E-05 |
| | cg23186534 | 2,34E-05 |
| | cg26386962 | 2,34E-05 |
| *SOX6* | cg08643003 | 3,90E-04 |
| | cg26830200 | 2,80E-02 |
| | cg00664860 | 4,04E-05 |
| | cg08224625 | 4,04E-05 |
| | cg09749669 | 4,04E-05 |

| | | |
|---|---|---|
| | cg10825530 | 4,04E-05 |
| | cg15497474 | 4,04E-05 |
| | cg21992400 | 4,04E-05 |
| | cg23530266 | 4,04E-05 |
| | cg24331727 | 4,04E-05 |
| *ZEB2* | cg08754046 | 1.65E-16 |
| | cg00025331 | 4.49E-04 |
| | cg03303633 | 4.49E-04 |
| | cg14807945 | 4.49E-04 |
| | cg16219453 | 3.00E-02 |
| | cg22267613 | 3.00E-02 |

**Table S7.** Bioinformatic tools, related to Star methods and supplemental computational methods.

| | | |
|---|---|---|
| Macs2 | (Feng et al., 2012) | https://pypi.org/project/MACS2/ https://github.com/macs3-project/MACS |
| refdata-cellranger-atac-hg38-version refdata- | | https://support.10xgenomics.com/single-cell-atac/software/downloads/ |
| JASPAR | (Bryne et al., 2008) | https://bioconductor.org/packages/release/data/annotation/html/JASPAR2020.html http://jaspar.genereg.net |
| Single Cell Experiment | (Amezquita et al., 2020) | https://bioconductor.org/packages/release/bioc/html/SingleCellExperiment.html |
| Escape | DOI:10.18129/B9.bioc.escape | https://github.com/ncborcherding/escape |
| ggseqlogo | | https://cran.r-project.org/web/packages/ggseqlogo/index.html |
| TFBSTools | | https://bioconductor.org/packages/release/bioc/html/TFBSTools.html |
| uwot | | https://cran.r-project.org/web/packages/uwot/index.html |
| viridisLite | (Garnier et al., 2021) | https://cran.r-project.org/web/packages/viridisLite/index.html |
| ShinyGO | (Ge et al., 2020) | http://bioinformatics.sdstate.edu/go/ |
| Python3 | (Van Rossum and Drake, 2009) | https://www.python.org/downloads/ |
| 10x Genomics Loupe Browser | | https://www.10xgenomics.com |

| | | |
|---|---|---|
| ComplexHeatmap | | https://jokergoo.github.io/ComplexHeatmap-reference/book/ |
| ggplot2 | (Wickham, 2009) | https://cran.r-project.org/web/packages/ggplot2/index.html |
| igraph | | https://igraph.org |
| IRanges | | https://bioconductor.org/packages/release/bioc/html/IRanges.html |
| reticulate | | https://cran.r-project.org/web/packages/reticulate/index.html |
| SummarizedExperiment | | https://bioconductor.org/packages/SummarizedExperiment. |
| tidyverse | (Wickham et al., 2019) | https://www.tidyverse.org/packages/ |
| ggpubr | (Kassambara, 2020) | https://cran.r-project.org/web/packages/ggpubr/index.html |
| SARTools | (Varet et al., 2016) | https://github.com/PF2-pasteur-fr/SARTools |
| pheatmap | | https://cran.r-project.org/web/packages/pheatmap/index.html |
| lluminaHumanMethylationEPICmanifest | | https://bioconductor.org/packages/release/data/annotation/html/IlluminaHumanMethylationEPICmanifest.html |
| IlluminaHumanMethylationEPICanno.ilm10b5.hg38 | | https://github.com/achilleasNP/IlluminaHumanMethylationEPICanno.ilm10b5.hg38 |
| Rstudio | | https://www.rstudio.com/ |

### Supplemental Computational Methods

#### Filtering and normalization of scRNA-seq data

The Seurat Package v.4.0. was used to perform quality control and normalization on the count matrices obtained after the Cellranger aggregate command. The gene count per cell, UMI count per cell and percentage of mitochondrial and ribosomal transcripts were computed using the functions of the Seurat package (Hao et al., 2021). Low quality cells expressing few genes (less than 200) were excluded from the downstream analysis. Genes expressed in less than three cells were removed. Measures of quality can also be based on the proportion of reads that map to genes in the mitochondrial genome. Cells with more mitochondria or more mitochondrial activity may naturally have larger mitochondrial proportions in the data. If the dataset is assumed to have high quality cells, outliers related to mitochondrial gene expression can be removed. The Median Absolute Deviations (MAD) is a numeric scalar giving the number of median absolute deviations to be used to flag potentially problematic cells based on total_counts (total number of counts for the cell, or library size) and total_features (number of features with non-zero expression) where cells are flagged for filtering only if total_features have nmads below the median (McCarthy et al., 2017). We used the MAD-based definition of outliers, using the Isoutlier function of scater package version 1.0.4, to remove putative low-quality cells from the dataset. Counts were adjusted for cell-specific sampling ('normalized') using the scTransform function with regression of cell cycle genes and mitochondrial content.

#### Cell-cycle state assessment using scRNA-seq data

The cell cycle state assessment was performed using a gene set and scoring system previously reported (Tirosh et al., 2016). The S and G2M scores were calculated based on 43 S phase-specific genes and 54 G2 or M phase-specific genes and the Seurat package cell cycle scoring function was used for calculation of actual scores. Manual evaluation of the assessment fits well with expected cell cycle state of the cells as the R1 cluster at Day 0 (unsynchronized hESCs) has the highest proportion of cells in S-phase, while the lowest percentage of G1-phase cells are found in cluster R12.

#### Clustering and Dimensionality reduction

The Resolution for finding clusters were computed using scClustviz SNN- based clustering. Then, principal component analysis was performed using RunPCA function of the Seurat package. For UMAP visualization, clusters were identified using FindNeighbors and FindClusters Seurat function using resolution 0.55, followed by the RunUMAP function across samples with the same parameters. UMAP preserves aspects of global structure in larger datasets and is therefore preferred for visualization over t-SNE (Becht et al., 2019). We used FindMarkers or FindAllMarkers functions to compute differentially expressed genes between the clusters. The UMAP plots and differentially expressed genes between clusters can be interactively browsed using the link provided in STAR methods software table.

#### CytoTRACE

We utilized CytoTRACE (Cellular (Cyto) Trajectory Reconstruction Analysis) with gene counts for all datasets (Days 0, 7, 13 and 20 merged, and Day 0 and Day 20 Merged) for prediction of differentiation state of cells from scRNA-seq data. In short, CytoTRACE leverages single cell gene counts, covariant gene expression and local neighbourhoods of transcriptionally similar cells, to predict ordered differentiation states from scRNA-seq data. CytoTRACE uses smoothing steps within the dataset to remove confounding factors associated with direct comparison of genes expressed by each cell in cross-study differences in depth and sensitivity.

**Single-cell ATAC sequencing analysis**

Cross platform linkage of scRNA-Seq data with a scATAC-seq data unconstrained and constrained integration aims to align cells from scATAC-seq and scRNA-seq experiments by combining them both together. A constrained integration can improve the quality of cross-platform alignment by limiting the alignment search space based on prior knowledge of the cell type. ArchR performs an unconstrained integration to determine preliminary cluster identities, followed by a refined constrained integration based on this prior knowledge (Granja et al., 2021). As part of the scATAC-seq and scRNA-seq integration, the GeneIntegration matrix containing linked gene expression data is added to HDF5 formatted arrow files. The GeneIntegrationMatrix enabled us to compare the linked gene expression with the gene score estimates of gene expression. The genes associated with known and new marker genes related to pluripotency, neuronal differentiation, and cell cycle of single cell RNA-seq were constrained and used as new labels of integrated clusters.

**Pseudo-Bulk replicates and Peak calling**

Using ArchR's addReproduciblePeakSet() function utilizing Macs2 for peak calling, we created a reproducible MACS2-derived merged peak set (Granja et al., 2021; Zhang et al., 2008). The new peak matrix was added to the ArchRproject using AddpeakMatrix function on MACS2-derived merged peak set. AddMarkerFeatures() was used to compute unique peaks for each cluster or group of clusters. Using the markerHeatmap() function, we visualized marker peaks in heatmaps, and we used the Plotbrowsertrack function to visualize the tracks. Motif enrichment was used to predict what transcription factors may mediate the binding events that create the accessible chromatin sites. By examining the differential peak opening in the different cell populations, we can identify motifs that have a high level of enrichment. A motif's presence in a peak set was determined using the AddmotifAnnotation function. PeakAnnotation was performed using peakAnnoEnrichment() on Jaspar2020 (Bryne et al., 2008). To compute representative seq logo for the motifs from position weight matrix we used the ggSeqlogo package (Wagih, 2017).

**ChromVAR Deviations Enrichment with ArchR**

ChromVAR deviations for each single cell were calculated using the “addDeviationsMatrix” function (Granja et al., 2021). To visualize transcription factor (TF) footprinting we used the “plotFootprints” function with the normalization method “subtract” which subtracts the Tn5 bias from the ATAC-seq footprint.

**TF Footprinting**

The integration of several levels of single cell data can provide new insights into gene regulation in cell populations. We used the ArchR addCoAccessibility() function to store the peak co-accessibility information and the getCoAccessibility function to retrieve it at the loop (Granja et al., 2021). Peak co-accessibility is a correlation in accessibility between two peaks across many single cells. We performed peak-coaccessibility analysis to predict regulatory interactions in the scATAC-seq dataset and in the integration of scATAC-seq and scRNA-seq data such that upstream enhancer activity can be predicted through peak-to-gene linkage analyses. We also used the Peak-to-gene linkage analysis to plot Peak2GeneHeatmap, containing two side-by-side heatmaps, one pertaining to scATAC-seq gene score data and the other to scRNA-seq gene expression data. Additionally, we analyzed the enrichment of TF binding motifs using the peakAnnoEnrichment function implemented in ArchR (Granja et al., 2021).

**Pseudo-time trajectory analysis**

We constructed a cellular trajectory spanning clusters R0 to R15, using the addTrajectory function implemented in ArchR on GeneScore, GeneIntegration, Peak and Tilematrix (Granja et al., 2021).

#### Glossary for webtools

Useful glossary information for functions in the web tools found at https://cancell.medisin.uio.no/ for easy access and exploration of the scRNA-seq and the chromatin accessibility data presented in this work.

**hESCNeuroDiff scRNA (https://cancell.medisin.uio.no/scrna/hescneurodiff/)**

| | |
|---|---|
| Orig.ident | Defined sample description |
| nCountRNA | Number of RNA counts in each cell |
| nFeatureRNA | Number of genes expressed in each cell |
| Percent.mt | Mitochondrial percentage in each cell |
| S.score | Cell cycle S phase analysis |
| G2M score | Cell cycle G2/M phase analysis |
| CC.difference | Regression score of cell cycle |
| Phase | Cell cycle phase analysis |
| Old.ident | Cluster number detected and named as default by Seurat with default threshold |
| ncount.SCT | Normalized count after SCTransform |
| nFeatureSCT | Normalized number of genes of SCTransform |
| SCT_snn_res.0.1 …0.9 | Cluster detection at resolutions 0.1 to 0.9 |
| Seurat_clusters | Cluster number detected and named as default by Seurat with the selected threshold |
| Original_seurat_clusters | Seurat clusters default settings |
| Pathway analysis | Pathway enriched terms |
| PC1/2/3/4/5 | Principal Component 1-5 |
| UMAP1/2 | Uniform Manifold Approximation and Projection for dimension reduction 1 or 2 |
| tSNE1/2 | t-distributed Stochastic Neighbour Embedding 1 or 2 |

**hESCNeuroDiff scATAC (https://cancell.medisin.uio.no/scatac/hescneurodiff.archr/)**

| | |
|---|---|
| Orig.ident | Sample names |
| Clusters | scATAC-seq clusters |
| Unconstrained | Unconstrained scATAC-seq clusters |
| Constrained | Integration of scATAC-seq clusters with original scRNA-seq clusters |
| Constrained remap | Integration of scATAC-seq with annotated scRNA-seq clusters |

| | |
|---|---|
| GeneScoreMatrix | scATAC-seq accessible peaks in vicinity of the gene based on custom ArchR distance-weighted accessibility models |
| GeneIntegrationMatrix | Integration of scATAC-seq with scRNA-seq, where accessible peaks in vicinity of selected gene is linked with the measured gene expression |
| PeakMatrix | Peak derived matrix of insertion counts of accessible regions |
| MotifMatrix | TF motif enrichment within accessible peaks |
| Co-accessibility | scATAC-seq correlations between two accessible peaks |
| Peak2GeneLinkage | Integration of scRNA-seq and scATAC-seq showing correlations between peak accessibility and expressed genes |
| Motif deviation score | highly correlated Transcription Factors expression with motif accessibility can be identified based on the correlation of the inferred gene score to the chromVAR motif deviation. Motif deviation score is computed per-cell deviations across all of motif annotations |
